# The SARS-CoV-2 accessory factor ORF7a downregulates MHC class I surface expression

**DOI:** 10.1101/2022.05.29.493850

**Authors:** Shuxuan Zheng, Hendrik de Buhr, Patrique Praest, Anouk Evers, Ingrid Brak-Boer, Mariëlle van Grinsven, Ylenia Longo, Liset de Vries, Wilco Nijenhuis, Lukas C. Kapitein, Jeffrey M. Beekman, Monique Nijhuis, Ingo Drexler, Emmanuel J. H. J. Wiertz, Robert Jan Lebbink

**Affiliations:** Department of Medical Microbiology, University Medical Center Utrecht, Utrecht, The Netherlands; Institute for Virology, Düsseldorf University Hospital, Heinrich-Heine-University, Düsseldorf, Germany; Cell Biology, Neurobiology and Biophysics, Department of Biology, Faculty of Science, Utrecht University, Utrecht, The Netherlands; Centre for Living Technologies, Alliance TU/e, WUR, UU, UMC Utrecht, Utrecht, The Netherlands; Department of Pediatric Pulmonology, Wilhelmina Children’s Hospital, University Medical Center Utrecht, Utrecht, The Netherlands; Regenerative Medicine Center Utrecht, University Medical Center, Utrecht, The Netherlands

**Author notes:** Correspondence (R.J. Lebbink). Equal contribution (Shuxuan Zheng, Hendrik de Buhr).

**Keywords:** SARS-CoV-2, ORF7a, MHC class I antigen presentation pathway, COVID-19, immune evasion

## Abstract

The pandemic caused by severe acute respiratory syndrome coronavirus 2 (SARS-CoV-2) has resulted in over 500 million infections and more than six million deaths worldwide. Although the viral genomes of SARS-CoV-1 and SARS-CoV-2 share high sequence homology, the clinical and pathological features of COVID-19 differ profoundly from those of SARS. It is apparent that changes in viral genes contribute to the increased transmissibility of SARS-CoV-2 and pathology of COVID-19.

Cytotoxic T lymphocytes play a key role in the elimination of virus-infected cells, mediated by recognition of virus-derived peptides that are presented on MHC class I molecules. Here, we show that SARS-CoV-2 can interfere with antigen presentation thereby evading immune surveillance. SARS-CoV-2 infection of monkey and human cell lines resulted in reduced cell-surface expression of MHC class I molecules. We identified a single viral gene product, the accessory factor open reading frame 7a (ORF7a), that mediates this effect. ORF7a interacts with HLA class I molecules in the ER, resulting in ER retention or impaired HLA heavy chain (HC) trafficking to the Golgi. Ultimately, these actions result in reduced HLA class I surface expression on infected cells. Whereas ORF7a from SARS-CoV-2 reduces surface HLA class I levels, the homologous ORF7a from the 2002 pandemic SARS-CoV-1 did not, suggesting that SARS-CoV-2 ORF7a acquired the ability to downregulate HLA-I during evolution of the virus. We identified a single amino acid in the SARS-CoV-1 ORF7a luminal domain that, upon mutating to the corresponding SARS-CoV-2 ORF7a sequence, induced a gain-of-function in HLA surface downregulation. By abrogating HLA class I antigen presentation via ORF7a, SARS-CoV-2 may evade host immune responses by inhibiting anti-viral cytotoxic T cell activity, thereby contributing to the pathology of COVID-19.

## Introduction

The severe acute respiratory syndrome coronavirus 2 (SARS-CoV-2) is the causative agent of the current coronavirus disease 2019 (COVID-19) pandemic. Whereas common circulating coronaviruses only induce mild illnesses in humans [1], pathology of COVID-19 is more severe. Over the past twenty years, two additional zoonotic coronaviruses have emerged that cause serious respiratory disease in humans: severe acute respiratory syndrome coronavirus (SARS-CoV-1) and Middle East respiratory syndrome coronavirus (MERS-CoV) [2]. Like SARS-CoV-1 and MERS-CoV, SARS-CoV-2 is an enveloped positive-sense single-stranded RNA virus from the family of *Coronaviridae* [3]. The SARS-CoV-2 genome consists of 14 open-reading frames (ORFs), that encode multiple proteins with dedicated functions. ORF1a and ORF1b encode 16 non-structural proteins (NSP1-NSP16) that comprise the replicase-transcriptase complex [4]. In addition, SARS-CoV-2 encodes 13 ORFs at the 3’ end of its genome, consisting of four structural and nine putative accessory factors. The structural proteins encode constituents of the viral nucleocapsid, including the spike (S), envelope (E), membrane (M) and nucleocapsid (N) proteins [5]. The nine accessory proteins are virulence factors involved in the pathogenicity of the virus [6]. Many of their specific functions remain unknown, although it is likely that they impact the host response to virus infection and as such contribute to the pathology of COVID-19.

The MHC class I antigen presentation pathway (in humans also referred to as the HLA class I antigen presentation pathway) is an integral component of the host’s immune defense to counteract viral infections. Viruses depend on their host to facilitate infection and replication, e.g. for viral protein synthesis. Upon viral protein synthesis, a fraction of these molecules are processed by the proteasome to small peptides that are subsequently translocated to the ER lumen by the transporter associated with antigen processing (TAP). There, they are loaded onto MHC class I molecules and presented at the cell surface to circulating CD8^+^ cytotoxic T lymphocytes (CTLs). Upon recognition of the MHC-Ι-peptide complex via their T cell receptor, CTLs become activated and induce killing of the antigen-presenting cells by various effector mechanisms [7]. To counteract this anti-viral activity, many viruses have evolved sophisticated mechanisms to interfere with antigen presentation [8]. They e.g. inhibit MHC class I molecule synthesis, induce their degradation via the proteasome, or avert peptide or MHC class I transport. These activities ultimately reduce MHC class I antigen presentation on the cell surface, thereby interfering with recognition by CTLs [8].

As CTLs play a key role in the elimination of virus-infected cells, we assessed whether SARS-CoV-2 has evolved strategies to counteract these antiviral measures by interfering with the HLA class I antigen presentation pathway. Indeed, our results suggest that the accessory factor open reading frame 7a (ORF7a) acts as an immune evasin by downregulating HLA class I molecules from the surface of infected cells. By abrogating HLA class I antigen presentation, ORF7a may facilitate evasion of cytotoxic T cell responses in infected individuals.

## Material and Methods

### Cell lines and viruses

HEK-293T and HK-1 cells were maintained in Roswell Park Memorial Institute medium (RPMI 1640; Life Technologies) supplemented with 5% FCS (Sigma), 2 mM L-glutamine, 100 U/mL penicillin, and 100 mg/mL streptomycin. The MelJuSo, Huh7 and A549-ACE2-TMPRSS2 cells were maintained in Dulbecco’s modified Eagle medium (DMEM; Life Technologies) supplemented with 5% FCS (Sigma), 2 mM L-glutamine, 100 U/mL penicillin, and 100 mg/mL streptomycin.

SARS-CoV-2 strain NL/2020 was obtained from the European Virus Archive global (EVAg, Ref-SKU 010V-03903). A ΔORF7a recombinant SARS-CoV-2 virus encoding the GFP gene at the ORF7a locus [9] and the parental SARS-CoV-2 isolate harbouring an intact ORF7a gene were kindly provided by Prof. Volker Thiel (Institute of Virology and Immunology, Bern, Switzerland). All viruses were propagated and titrated on Vero E6 cells using the tissue culture infective dose 50 (TCID50) endpoint dilution method. Propagation and experiments with infectious SARS-CoV-2 viruses were conducted in our BSL3 lab.

### Plasmids

The plasmids of the SARS-CoV-2 cDNA library as cloned in the pLVX-EF1alpha-IRES-Puro (Takara/Clontech) vector were a kind gift from Prof. Nevan Krogan (University of California San Francisco, USA) [10]. The library lacked NSP3 and NSP16. Additionally, as the viral Spike gene was provided in a different backbone vector, the cDNA was transferred to pLVX-EF1alpha-IRES-Puro (Takara/Clontech) by classic restriction digest and ligation. Lentiviral cDNA plasmids were transduced into HEK-293T cells and after antibiotic selection subjected to HLA class I surface level quantification by flow cytometry. Cell lines expressing NSP1 and NSP14 did not survive transduction and subsequent antibiotic selection and were, therefore, excluded from the analysis For follow-up studies, we cloned a T2A-mAmetrine cassette in frame downstream of the PuroR gene in the pLV-CMV-IRES-PuroR vector. This vector (pLV-CMV-IRES-PuroR-T2A-mAmetrine; a.k.a. RP-1023) served as backbone vector for all subsequent cDNA variants. We cloned untagged and C-terminally StrepII-tagged variants of SARS-CoV-2 ORF7a, SARS-CoV-2 ORF8, and SARS-CoV-1 ORF7a Gibson cloning procedures in the RP-1023 vector. For SARS-CoV-1 ORF7a, overlapping ±90 nt ssDNA oligos (IDT Europe) were assembled by overlap extension PCR and cloned (with and without a C-terminal Strep-II tag) in the RP-1023 vector. The sequences of the coding regions of ORF7a and ORF8 (both non-tagged and StrepII-tagged) are presented in Table S1). Chimeric ORF7a molecules, comprising of indicated sequences from both SARS-CoV-1 ORF7a and SARS-CoV-2 ORF7a were constructed by PCR amplification using primers that incorporated the desired nucleotide substitutions (primers used are presented in Table S2). All vectors were sequence-verified by Sanger sequencing.

ACE2-GFP and TMPRSS2-HA for the A549-ACE2-TMPRSS2 cells were cloned from vectors kindly provided by Dr. Wentao Li and Dr. Berend Jan Bosch (Utrecht University, Utrecht, The Netherlands) into lentiviral vectors, where they were expressed from a hEF1A promoter. The lentiviral plasmid also contains a PGK promoter driving expression of a blasticidin resistance gene and a BlastR-T2A-eGFP cassette for the ACE2-GFP and TMPRSS2-HA expression vector respectively.

### Lentivirus transductions

For the pLVX-EF1alpha-IRES-Puro vector system, lentiviral particles were prepared using the Lenti-X™ Packaging system (Takara Bio). For the pLV-CMV-IRES-PuroR-T2A-mAmetrine vectors, lentiviruses were produced using standard lentiviral production protocols with third-generation packaging vectors. Upon lentivirus production, viral supernatants were centrifuged at 3000rpm and stored at −80°C until further use. In general, ∼25k target cells were infected with ∼100 µl lentivirus supernatants in 24 well plates.

### SARS-CoV-2 infections

SARS-CoV-2 viruses (see section ‘cell lines and viruses’) propagated in Vero E6 cells were used to infect Vero E6 or A549-ACE2-TMPRSS2 cells. Briefly, cells were plated in a 24 well plate the day before infection, and subsequently mock infected or infected at a multiplicity of infection (MOI) of 0.01 (for Vero E6 cells) or 0.5 (for A549-ACE2-TMPRSS2 cells) in DMEM supplemented with 2% FCS. After a 1-hour incubation, the inoculum was removed and cells were maintained in complete medium for one or two days. Subsequently, cells were detached by TrypLE™ (Thermo) and fixed in 4% PFA prior to surface MHC-I and Spike antibody stainings. Cells were subjected to flow cytometry (BD FACS Canto II) and the data was analyzed with FlowJo (BD Biosciences) software.

### Intracellular staining

Cell lines used for intracellular HLA-I stainings were harvested using trypsin (Gibco) and pelleted by centrifugation. The supernatant was removed, cells were washed in icecold PBA (PBS + 0.5% BSA + 0.02% NaAzide) and pelleted again. Cells were then fixed in icecold 3.7% PFA in PBS for 10 min. After a washing step with icecold PBS, cells were incubated in icecold PBS + 0.5% saponin + 2% FCS for 10 min. Cells were then incubated for 30 min at 4°C with the conjugated antibody in icecold PBS + 0.5% saponin + 2% FCS and after two subsequent washing steps fixed with icecold 3.7% PFA in PBA and subjected to flow cytometry analysis (BD FACS Canto II). The antibody used was: PE-conjugated W6/32 (Serotec MCA81PE, 1:5) [11].

### Surface staining

Cell lines used for surface stainings were harvested using trypsin (Gibco) and pelleted by centrifugation. The supernatant was removed, cells were washed in icecold PBA (PBS + 0.5% BSA + 0.02% NaAzide) and pelleted again. Cells were then incubated for 30 min at 4°C with the directly-conjugated antibody and after two subsequent washing steps fixed with icecold 3.7% PFA in PBA and subjected to flow cytometry analysis (BD FACS Canto II). The antibodies used were: PE-conjugated W6/32 (Serotec MCA81PE, 1:10) [11], anti-CD9-FITC (BD Pharmingen 555371, 1:40), anti-CD46-FITC (BD Pharmingen 555949, 1:20), anti-CD49b-PE (BD Pharmingen 555669, 1:20), anti-CD58-PE (BD Pharmingen 555921, 1:30), anti-CDw119-PE (BD Pharmingen 558934, 1:20).

The primary antibody used for Spike staining was human anti-SARS-CoV-2 Spike (1:682, REGN #10987), which was kindly provided by Wentao Li and Berend Jan Bosch (Utrecht University, Utrecht, The Netherlands). The goat anti-human IgM+ IgG (H+L) (1:160, Jackson #109-116-127) antibody was used as secondary antibody

### Inhibition of proteasome and p97

For proteasome inhibition, we employed 20 μM MG132 (Sigma-Aldrich, Zwijndrecht, NL, C2211-5MG) and for p97 inhibition 4 μM CB-5083 (HY-12861; MCE) for 4h each.

### Immunoblotting and immunoprecipitations

Following cell lysis in Triton (1% Triton X-100, 50 mM Tris HCl [pH 7.5], 150 mM NaCl) lysis buffer supplemented with 10 μM Leupeptin (Roche), and 1 mM AEBSF, proteins were denatured in reducing sample buffer. Samples were incubated at 95°C for 5 min prior to loading. For SDS-PAGE, 4-12% Bolt Bis-Tris Plus SDS gels (Thermo Scientific) were used. The proteins were transferred to Trans-Blot Turbo PVDF membranes (BIO-RAD) using a Trans-Blot Turbo transfer system (BIO-RAD) for 10 minutes at 25V. Membranes were incubated for 1h at room temperature (RT) in PBS containing 0.05% Tween 20 (PBST) and 4% milk powder (Campina). Primary antibodies were diluted in PBST supplemented with 1% milk powder and incubated overnight at 4°C. Membranes were washed three times for 5 min with PBST at RT. HRP-labelled secondary antibodies were diluted in PBST supplemented with 1% milk powder and used to incubate the membranes in the dark for two hours at 4°C, or one hour at RT. Membranes were again washed three times for 5 min with PBST at RT before detection. Pierce ECL Western Blotting Substrate (Thermo Scientific) was added to the membranes for detection of chemiluminescence, followed by image acquisition (Image Quant LAS 4000). The primary antibodies used for immunoblotting were: HC10, monoclonal mouse anti-SARS-CoV-2 ORF7a (Genetex, 632602, 1:1000), polyclonal rabbit anti-SARS-CoV-2 ORF8a (Genetex, 135591, 1:1000), monoclonal transferrin receptor antibody (H68.4, Invitrogen, 1:1000) and monoclonal StrepII (C23.21, purified in our lab). Secondary antibodies used were goat anti-mouse IgG-HRP (115-035-174, Jackson ImmunoResearch Europe Ltd, 1:10000) and mouse anti-rabbit IgG-HRP (211-032-171, Jackson ImmunoResearch Europe Ltd, 1:10000). For empty vector and ORF7a immunoprecipitation 30-40 million cells per condition were taken up in Triton (1% Triton X-100, 50 mM Tris HCl [pH 7.5], 150 mM NaCl) lysis buffer supplemented with 10 μM Leupeptin (Roche), and 1 mM AEBSF at 7-10 days post transduction. For StrepII IP 25 μl Streptactin Sepharose® High Performance beads (GE Healthcare, GE28-9355-99) and for HLA-I IP 25 μl Protein G Sepharose® 4 Fast Flow (GE Healthcare, GE17-0618-01) were employed with the HC10 antibody for HLA-IP overnight at 4°C. Glycosylation status was detected by treatment with Endo H_f_ (P0703L, NEB) and N-Glycosidase F (11365177001, Roche) for 1h at 37°C prior to loading for SDS-PAGE.

### Immunofluorescence

Cells were fixed in 4% paraformaldehyde and 4% sucrose in PBS for 10 min with prewarmed fixative. After fixation, cells were washed in PBS, permeabilized in 0.5% Triton X-100 in PBS for 10 min and blocked in 3% BSA in PBS for 15 min, incubated with primary antibodies for 16 h at 4°C, washed three times with 0.1% Triton X-100 in PBS (PBST) for 10 min, and incubated with secondary antibodies together with DAPI for 2h at room temperature. Coverslips were then washed 2x with PBST and 1x with PBS each for 10 min, rinsed with 96% ethanol, dried, and mounted using antifade (ProLong Diamond; Molecular Probes). The following antibodies were used: mouse anti-SARS-CoV-2 ORF7a (Genetex, 632602, 1:1000) and mouse anti-W6/32 (own production from hybridoma, 1:1000). Secondary antibodies used were goat anti-mouse IgG2a cross-adsorbed secondary antibody, Alexa Fluor 594 (Thermo Fisher, A-21135, 1:600) and goat anti-mouse IgG1 cross-adsorbed secondary antibody, Alexa Fluor 647 (Thermo Fisher, A-21240, 1:600), together with DAPI (Sigma-Aldrich, 1:1000).

### Microscopy

Images were acquired using a Zeiss LSM880 Fast Airyscan microscope with 405 nm, 561 nm, 633 nm and argon multiline lasers, internal Zeiss spectral 3 PMT detector and spectroscopic detection using a Plan-Apochromat 63x/1.2 glycerol objective.

### Protein sequence alignments

Protein sequence alignments were made using MUSCLE in the Snapgene software package (GSL Biotech). ORF7a from the following virus isolates were aligned: SARS-CoV-2 (QZX66581.1), SARS-CoV-1 (SARS coronavirus Tor2; YP_009825057.1), Pangolin coronavirus (QIG55950.1), Bat coronavirus BM48-31 (ADK66846.1), Bat SARS coronavirus HKU3-2 (AAZ41334.1), Bat Betacoronavirus sp. RmYN07 (QWN56217.1), Bat Betacoronavirus sp. RpYN06 (QWN56257.1), and Bat Betacoronavirus sp. RsYN03 (QWN56237.1).

## Results

### MHC class I surface expression is downregulated upon SARS-CoV-2 Infection

The MHC class I antigen presentation pathway is a common target for viruses to evade the host’s immune system. We asked whether also SARS-CoV-2 can interfere with this cellular defense mechanism. We infected Vero E6 cells with the SARS-CoV-2 NL/2020 strain at an MOI of 0.01 and assessed MHC class I surface levels by flow cytometry at 1 and 2 dpi. Indeed, MHC class I surface levels were downregulated in Spike-expressing cells as compared to uninfected cells present in the same culture (Fig 1A) at 2 dpi, but not at 1 dpi. We next measured the effect of SARS-CoV-2 infection in human cells, for which we transduced A549 cells with ACE2 and TMPRSS2 and subsequently challenged these with SARS-CoV-2 NL/2020 at an MOI of 0.5. Also here, HLA class I surface levels were reduced in Spike-expressing cells at 2 dpi (Fig 1B). Our results show that SARS-CoV-2 infection induces downregulation of MHC class I surface levels in both a monkey and a human cell line.

**Figure 1.**
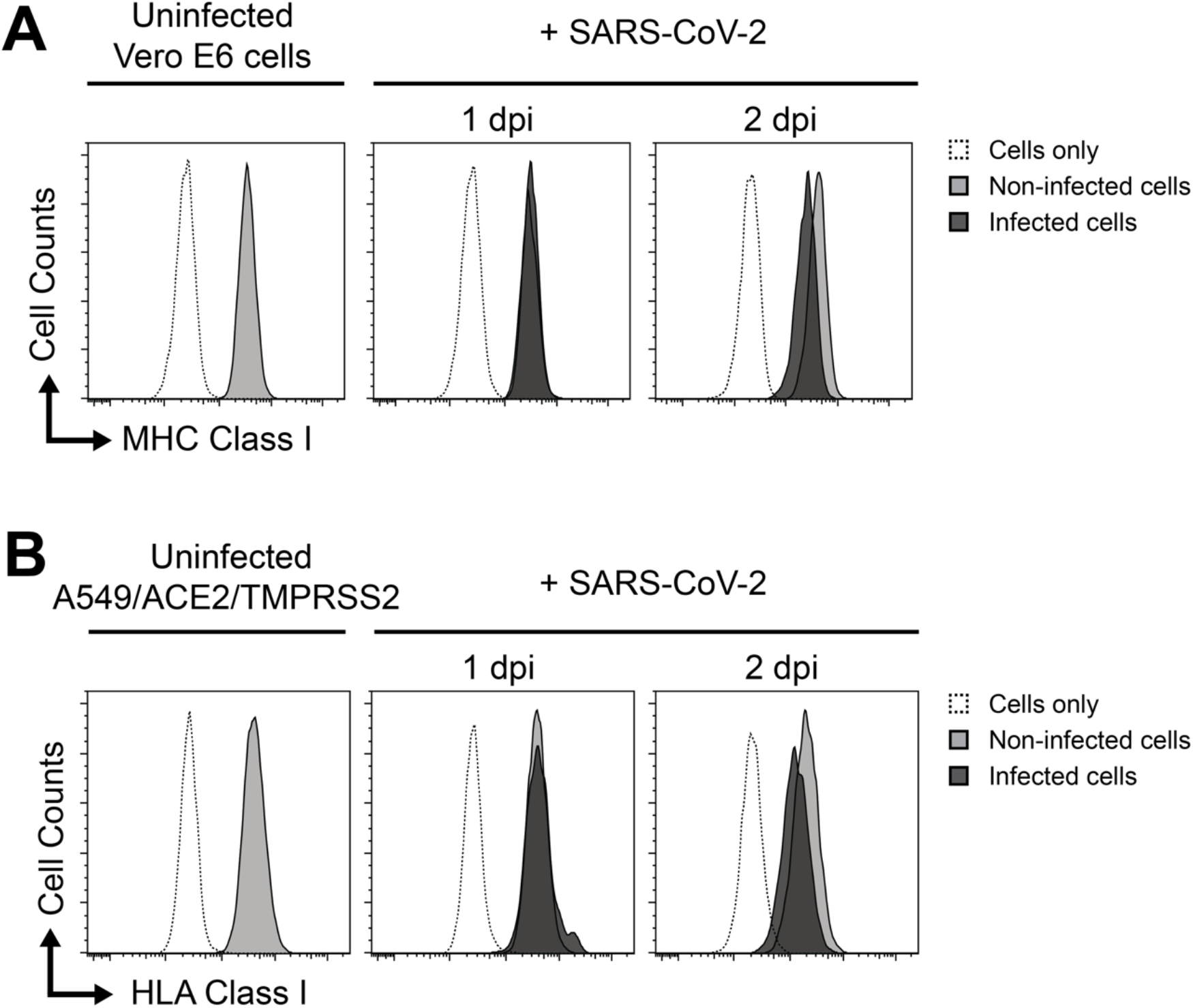
SARS-CoV-2 infection causes downregulation of surface MHC class I levels in Vero E6 and A549 cells. (**A**) Vero E6 cells were infected with SARS-CoV-2 strain NL/2020 (MOI 0.01) and analyzed for surface MHC class I levels by flow cytometry at 1 and 2 dpi. Spike antibody stains were included to allow discrimination between SARS-CoV-2 infected (infected, black filled histograms) and non-infected cells present in the same culture (gray histograms). Unstained cells are indicated (open histograms). (**B**) same as in A), yet A549-ACE2-TMPRSS2 cells were infected with SARS-CoV-2 strain NL/2020 at an MOI of 0.5. For human cells HLA-I nomenclature was used instead of MHC-I

### SARS-CoV-2 ORF7a downregulates surface HLA-I

The SARS-CoV-2 genome consists of 14 open-reading frames (ORFs), which encode for 31 viral proteins [12]. To assess which viral gene product causes surface HLA class I downregulation, we transduced a library of StrepII-tagged SARS-CoV-2 cDNAs in 293T cells and monitored HLA-I surface levels by flow cytometry (Fig 2A). Of all viral cDNAs, only ORF7a showed a significant downregulation of surface HLA-I levels of ±55% as compared to wt control cells (Fig 2B). Unexpectedly, ORF8, which was previously identified to downregulate HLA class I levels [13], did not impact HLA class I surface levels (Fig 2B).

**Figure 2.**
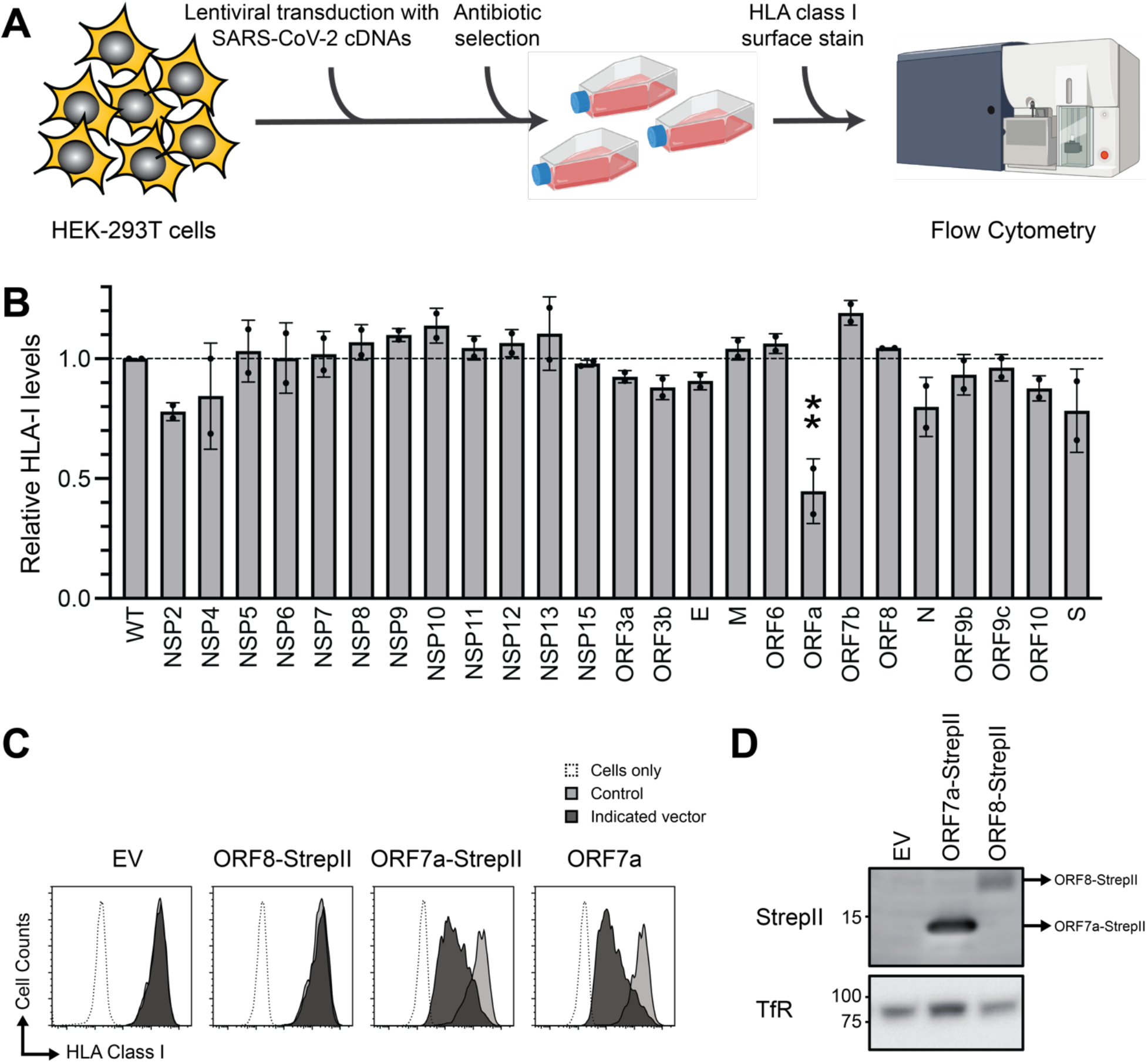
SARS-CoV-2 ORF7a causes downregulation of cell surface HLA class I levels. (**A**) Schematic overview of the screen to assess which SARS-CoV-2 cDNAs causes downregulation of cell surface HLA-class I levels in 293T cells. **(B)** Indicated SARS-CoV-2 cDNAs were introduced in 293T cells by lentiviral transduction and selected to purity. HLA class I protein surface levels were subsequently assessed by flow cytometry using an anti-HLA-I antibody (W6/32). Relative surface HLA-I levels as compared to non-transduced cells (wt) are presented. One-way ANOVA with Tukey’s multiple comparisons test for n = 2. P value ORF7a = 0,0011, rest not significant. (**C**) Flow cytometry analysis of surface HLA-I expression levels of 293T cells transduced with lentiviral vectors encoding SARS-CoV-2 ORF7a, ORF7a-StrepII, ORF8-StrepII, or empty vector control (EV) at 7 dpi. The cDNA vectors co-express the mAmetrine-fluorescent protein to allow discrimination between transduced (mAmetrine-positive) and non-transduced (mAmetrine-negative) cells within the same culture. Unstained cells are indicated (open histograms). One representative figure of >5 experiments is shown. (**D**) Western blot analysis of StrepII-tagged proteins from C): empty control vector (EV), SARS-CoV-2 ORF7a-StrepII or SARS-CoV-2 ORF8-StrepII. Transferrin receptor (TfR) protein levels are presented as loading control. One representative experiment of two is shown.

To validate our finding, we subcloned the ORF7a cDNA in a different lentiviral vector. Also here, ORF7a constructs potently downregulated HLA class I from the cell surface, whereas the ORF8 cDNA did not (Fig 2C). Both untagged and StrepII-tagged ORF7a induced potent downregulation of surface HLA class I (Fig 2C), whereas neither untagged nor StrepII-tagged ORF8 reduced surface HLA class I levels (Fig 2C and data not shown). The lack of HLA-I downregulation by ORF8 could potentially be explained by a relatively low protein expression of ORF8 (Fig 2D), which was apparent by Western blotting for the StrepII-tag located on the C-terminus of the protein. When we followed HLA-I surface expression in ORF7a-transduced cells in time, we observed that the HLA-I downregulation was transient and variable, and the phenotype was eventually lost after approximately two weeks (data not shown). Our results show that ORF7a, but not ORF8, downregulates HLA class I protein levels from the cell surface.

### ORF7a downregulates surface HLA-I in multiple cell lines and is specific to HLA class I

SARS-CoV-2 ORF7a induced surface downregulation of HLA-I expression. To assess whether ORF7a specifically downregulates HLA-I from the cell surface or rather impacts expression of proteins that migrate through the secretory pathway, we monitored the surface expression of multiple cell surface antigens in ORF7a-transduced 293T cells and compared this to the levels of these antigens in cells transduced with the other SARS-CoV-2 cDNAs (Fig 3A). Besides HLA class I, ORF7a did not impact the expression of CD9, CD46, CD49b, CD58 and CDw119 from 293T cells (Fig 3A and 3B). This implies that ORF7a does not cause a general defect in the secretory pathway that could explain for the loss of HLA-I from the cell surface, but rather that ORF7a specifically induces HLA-I surface downregulation.

**Figure 3.**
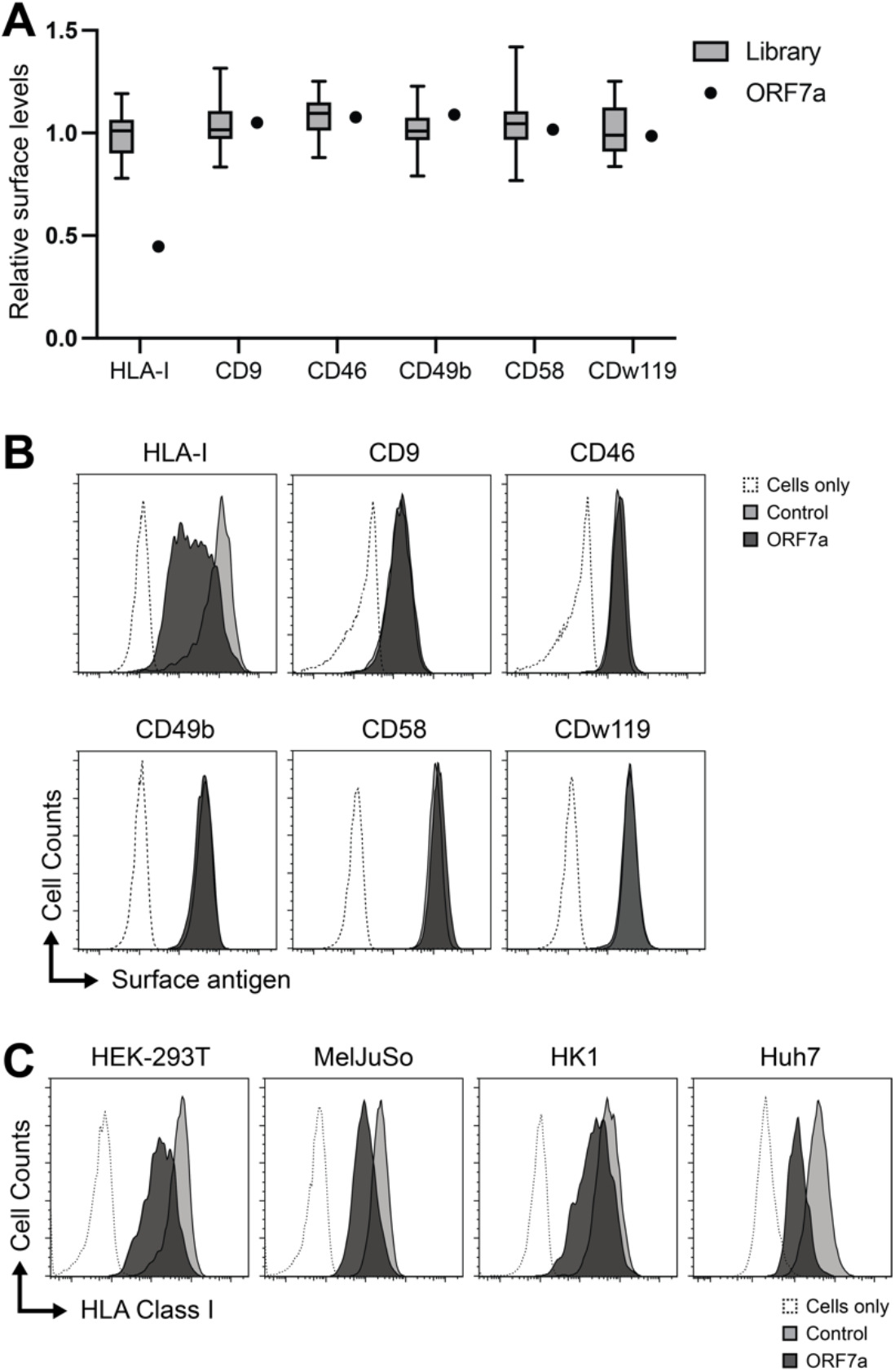
ORF7a specifically downregulates HLA-I from the cell surface of multiple cell types. (**A**) 293T cells transduced with StrepII-tagged SARS-CoV-2 cDNAs and subsequently selected to purity were stained with monoclonal antibodies to detect the cell-surface expression levels of the indicated proteins. To assess whether ORF7a-transduced cells showed differential cell-surface expression levels as compared to cells transduced with other (non-ORF7a) SARS-CoV-2 cDNAs, we pooled all non-ORF7a-cDNA-transduced cells in the analysis (indicated as ‘Library’) and compared these with the ORF7a-transduced cells. Expression levels for each antigen were normalized to stainings performed on non-transduced cells. (**B**) Flow cytometry analysis of surface expression levels of the indicated antigens in 293T cells transduced with StrepII-tagged SARS-CoV-2 ORF7a (determined by mAmetrine expression, dark gray histograms) compared to non-transduced cells (light gray histograms). Unstained cells are indicated (open histograms). Cells were analyzed at 4 dpi. (**C**) SARS-CoV-2 ORF7a downregulates HLA-I from the surface of multiple cell types. Flow cytometry analysis of surface HLA-I levels in 293T, MelJuSo, HK-1 and Huh7 cells transduced with a StrepII-tagged SARS-CoV-2 ORF7a (determined by mAmetrine expression, dark gray histograms) compared to non-transduced cells (light gray histograms). Unstained cells are indicated (open histograms). Cells were analyzed at 6 dpi. Note that for Huh7 cells, gating on mAmetrine was not possible due to relatively low expression of the fluorescent protein, therefore puromycin-selected cells (dark gray histogram) were compared with stained non-transduced cells (light gray histogram). One representative experiment of three experiments is shown

Next, we set out to confirm whether ORF7a also regulates HLA-I in additional cell lines. For this we transduced ORF7a in the cutaneous melanoma cell line MelJuSo, the nasopharyngeal carcinoma cell line HK-1 and the adult hepatocellular carcinoma cell line Huh7 (Fig. 3C). As expected, all these ORF7a expressing cell lines displayed reduced HLA-I surface expression levels, although the magnitude of downregulation varied between the cell lines.

We conclude that SARS-CoV-2 ORF7a specifically downregulates HLA-I from the surface of multiple human cell lines

### The ORF7a signal peptide sequence and transmembrane domain are essential for HLA-I surface downregulation

SARS-CoV-2 ORF7a is an 121 aa ER-resident protein consisting of a 15 aa signal peptide sequence, followed by an 81 aa luminal domain, a 20 aa transmembrane (TM) domain and a 5 aa tail [12]. To pinpoint important domains of ORF7a that are crucial for HLA-I downregulation, we generated different SARS-CoV-2 mutants (Fig. 4A) and assessed whether HLA-I surface regulation was impaired.

**Figure 4.**
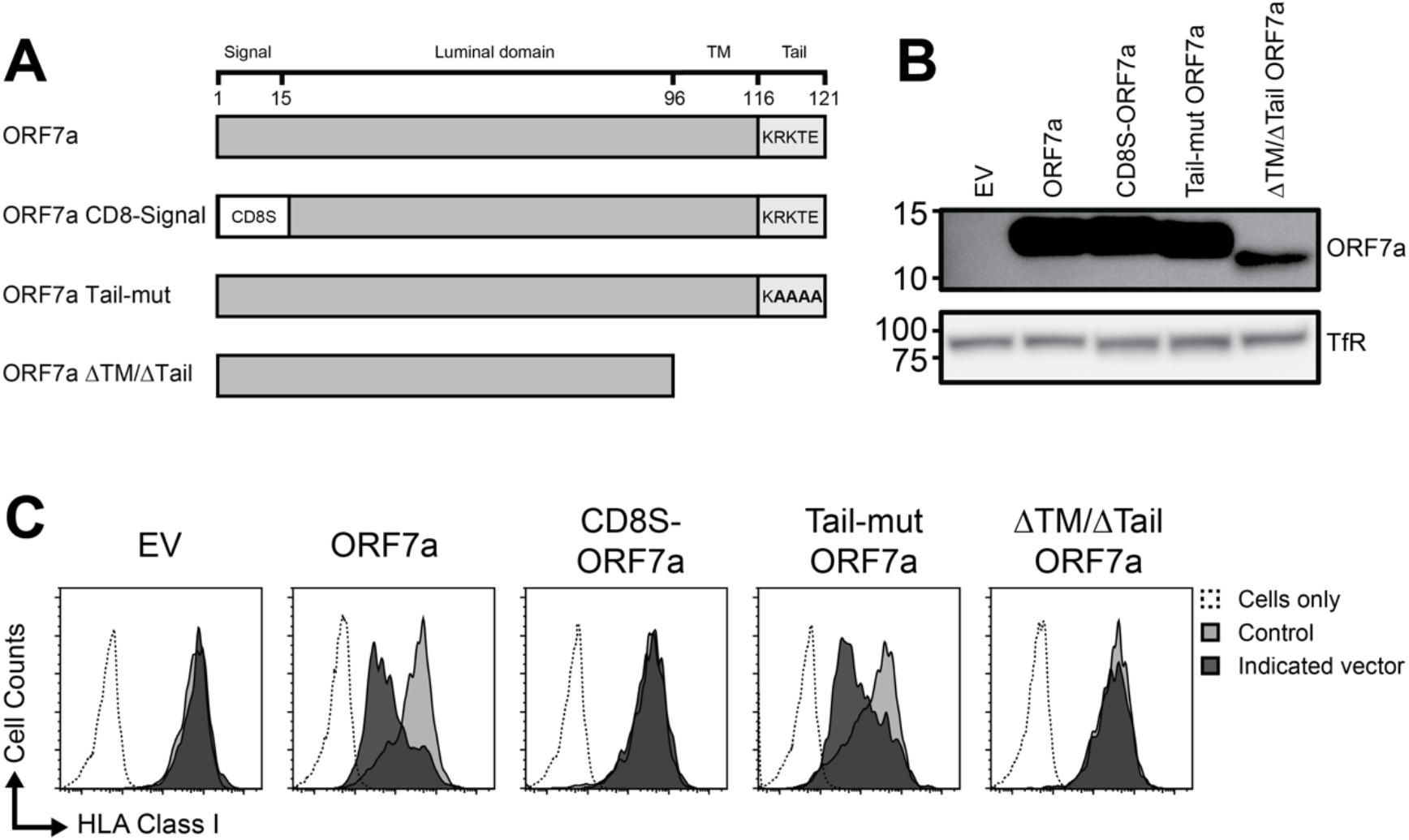
The ORF7a signal peptide sequence and transmembrane domain are essential for HLA-I surface downregulation, whereas the C-terminal ER-retrieval signal is not. (**A**) Schematic representation of the SARS-CoV-2 ORF7a protein, consisting of the signal peptide sequence, luminal domain, transmembrane (TM) domain and cytosolic tail, and the generated mutants CD8S-ORF7a, ORF7a Tail-mut and ORF7a ΔTM/tail. (**B**) 293T cells transduced with the ORF7a mutants from A) were selected by puromycin-treatment and expression levels were assessed by immunoblotting using an anti-ORF7a antibody and transferrin receptor antibody (TfR) as control. (**C**) 293T cells transduced with SARS-CoV-2 ORF7a, CD8S-ORF7a, ORF7a Tail-mut, ORF7a ΔTM/tail, or control vector (EV) were subjected to flow cytometry at 7 dpi to assess HLA-I cell-surface levels. mAmetrine expression was used to allow for discrimination between transduced (closed dark gray histograms, mAmetrine-positive) and non-transduced (light gray histograms, mAmetrine-negative) cells within the same culture. Unstained cells are indicated (open histograms). One representative experiment of five is presented.

We generated three mutants, replacing the ORF7a signal peptide sequence with the signal peptide sequence of human CD8A (CD8S-ORF7a), mutating the extreme C-terminal endoplasmic reticulum (ER) retrieval motif KRKTE [14] to KAAAA (ORF7a Tail-mut), and deleting the transmembrane and cytoplasmic tail completely (ORF7a ΔTM/Δtail, Fig. 4A). The different ORF7a variants were expressed in 293T cells (Fig 4B) and cell-surface expression of HLA-I was assessed by flow cytometry at 6 dpi (Fig. 4C). As seen previously, ORF7a expression induced potent surface HLA-I downregulation. Similar downregulation was observed upon introduction of the ORF7a tail-mut variant in which the ER retrieval motif was mutated. On the contrary, both the CD8S-ORF7a and the ORF7a ΔTM/Δtail mutant lost the ability to downregulate HLA-I surface expression. Our results suggest that presence of the ORF7a signal peptide sequence and transmembrane domain are crucial for HLA-I downregulation, whereas the ER-retrieval motif is dispensable for this activity.

### A single amino acid substitution of SARS-CoV-1 ORF7a confers the ability to downregulate surface HLA-I expression

SARS-CoV-2 ORF7a shares high sequence homology with SARS-CoV-1 ORF7a (multiple sequence alignment likelihood of 0,83) [15]. However, introduction of the SARS-CoV-1 ORF7a protein from the 2002 outbreak in 293T cells did not result in HLA-I surface downregulation (Fig 5C, upper right histogram). Therefore, HLA-I regulation appeared to be acquired by SARS-CoV-2 ORF7a upon evolution of the SARS-CoV-1 ORF7a gene. Protein-sequence alignment of both ORF7a proteins indicated that the sequence differences were predominantly located in three distinct ‘regions’ in the protein: in the signal sequence, a small area in the luminal domain, and the stalk-TM region (Fig. 5A). We reasoned that individual or multiple of these amino acid differences could be responsible for the gain-of-function phenotype observed in SARS-CoV-2 ORF7a. To study this, we replaced the three regions harbouring amino acid substitution of SARS-CoV-1 ORF7a with the corresponding sequence of SARS-CoV-2 ORF7a (Fig. 5B) and assessed whether the chimeric SARS-CoV-1 ORF7a proteins gained the ability to downregulate surface HLA-I expression. Indeed, whereas the wt SARS-CoV-1 ORF7a and two of the chimeric proteins (I and III) did not downregulate HLA-I, transfer of the luminal ‘region II’ from SARS-CoV-2 ORF7a to SARS-CoV-1 ORF7a resulted in potent downregulation of HLA-I from the cell surface (Fig. 5C). The transferred region II of SARS-CoV-2 only has 6 aa differences as compared to the same region of SARS-CoV-1 ORF7a. To identify which aa changes were responsible for the gain-of-function phenotype, we generated two chimeric mutants in which three amino acid changes were introduced in the SARS-CoV-1 ORF7a sequence. Whereas one of these chimeric ORF7a molecules (T71V/R72K/T74V) was not able to downregulate surface HLA-I levels (Fig. 5E, second overlay), introduction of the T59F/H62Q/A68P chimeric ORF7a molecule in 293T cells resulted in strong HLA-I surface downregulation (Fig. 5E). We next introduced all possible double and single aa changes from these three amino acids into the SARS-CoV-1 ORF7a molecule and noticed that those SARS-CoV-1 mutants that carried the T59F substitution gained the ability to downregulate HLA-I surface levels (Fig. 5E). Also, the H62Q substitution induced minor HLA-I surface downregulation, whereas the A68P substitution did not (Fig. 5E, right two overlays). We conclude that a single amino acid substitution in SARS-CoV-1 ORF7a (T59F) caused the gain-of-function phenotype in which surface HLA class I downregulation was acquired.

**Figure 5.**
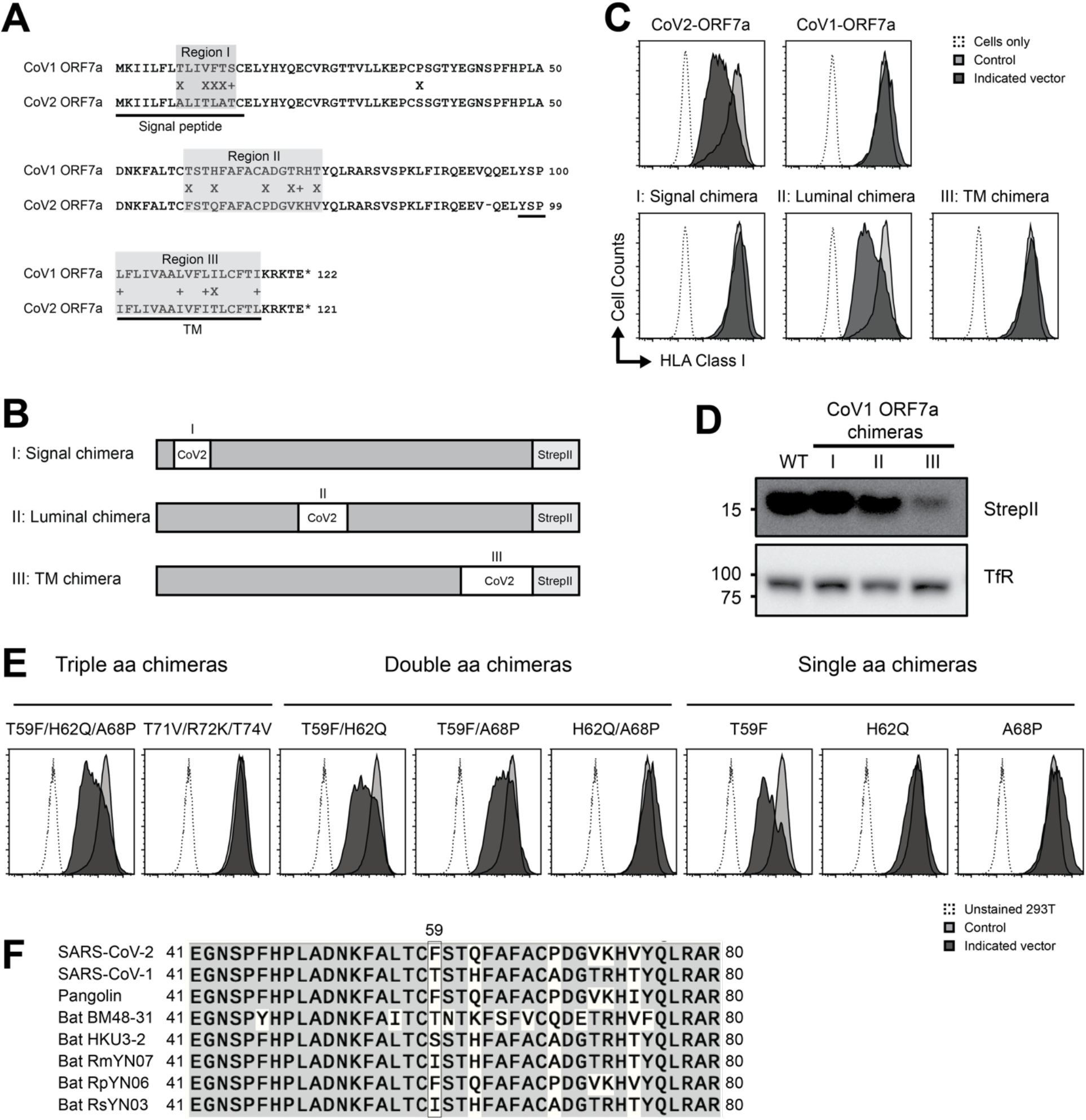
A single amino acid substitution in SARS-CoV-1 ORF7a confers the ability to downregulate surface HLA-I expression. (**A**) Protein sequence alignment of ORF7a from SARS-CoV-2 and SARS-CoV-1 identifies three regions of variability located in the signal peptide region (I), the luminal domain (II), and the stalk/TM region (III). The location of the signal peptide and TM region is indicated. (**B**) Schematic representation of the signal peptide chimera, the luminal chimera and the TM chimera, in which parts of the SARS-CoV-2 ORF7a sequence (indicated as open boxes) were introduced in the SARS-CoV-1 ORF7a sequence (gray boxes). (**C**) 293T cells transduced with the ORF7a chimeras from B) were subjected to flow cytometry at 6 dpi to assess HLA-I cell-surface levels. The ORF7a vector co-expresses mAmetrine to allow discrimination between ORF7a-transduced (closed black histograms, mAmetrine-positive) and non-transduced (open histograms, mAmetrine-negative) cells within the same culture. Unstained cells are indicated (open histograms). One representative experiment of three is presented. (**D**) ORF7a chimera and transferrin protein levels were assessed by immunoblotting of lysates from 293T cells expressing SARS-CoV-1-ORF7a, the signal peptide chimera, the luminal chimera or the TM chimera from C). (**E**) Indicated triple, double, and single aa mutations were introduced in the SARS-CoV-1 ORF7a protein. 293T cells were subsequently transduced with the chimeric molecules and subjected to flow cytometry at 6 dpi to assess HLA-I cell-surface levels, as described above. Unstained 293T cells are indicated (open histograms). (**F**) Protein sequence alignment of multiple isolates from human (SARS-CoV-1 and SARS-CoV-2), pangolin, and bat sarbecoviruses. Identical residues are indicated in gray. The amino acids at position 59 are indicated.

To assess whether the phenylalanine at position 59 in the SARS-CoV-2 ORF7a is also present in other sarbecovirus isolates, we performed a protein-sequence alignment of multiple ORF7a sequences obtained from the NCBI genome database (Fig. 5F). Indeed, we could readily identify sarbecovirus isolates of pangolin and bat origin carrying this specific amino acid. It is conceivable that these coronaviruses can also regulate surface MHC class I expression upon infection of cells in their host, although we have not addressed this. It was apparent that position 59 of ORF7a has considerate variability among virus isolates (Fig. 5F).

SARS-CoV-1 ORF7a has been shown to induce apoptosis in infected target cells [16–18]. Indeed, 293T cells transduced with SARS-CoV-1 ORF7a were quickly lost from the population of cells within ± one week after transduction, whereas SARS-CoV-2 ORF7a-transduced cells were not (data not shown). Therefore, the gain-of-function in HLA-I downregulation of SARS-CoV-2 ORF7a is accompanied by a loss-of-function in its ability to induce apoptosis. All chimeric SARS-CoV-1 ORF7a mutants we constructed induced apoptosis (data not shown), suggesting that the amino acids substitution studied here are not essential for SARS-CoV-1 ORF7a-mediated apoptosis induction.

### SARS-CoV-2 ORF7a induces intracellular HLA-I accumulation predominantly located in the ER

To assess whether ORF7a induces surface HLA class I downregulation by protein degradation, we assessed whether the total pool (both intracellular and surface expression) of HLA-I is depleted upon ORF7a expression in 293T cells. Unexpectedly, intracellular levels of HLA class I were potently upregulated in ORF7a-expressing 293T cells at 7 dpi, whereas HLA class I surface expression was downregulated (Fig. 6A) as observed previously. Western blot analysis confirmed that HLA-class I levels were upregulated in ORF7a-expressing cells as compared to EV controls (Fig 6B). Also, intracellular accumulation of HLA-I persisted in ORF7a expressing cells, even at late timepoints (21 dpi) when surface downregulation was already lost (data not shown).

**Figure 6.**
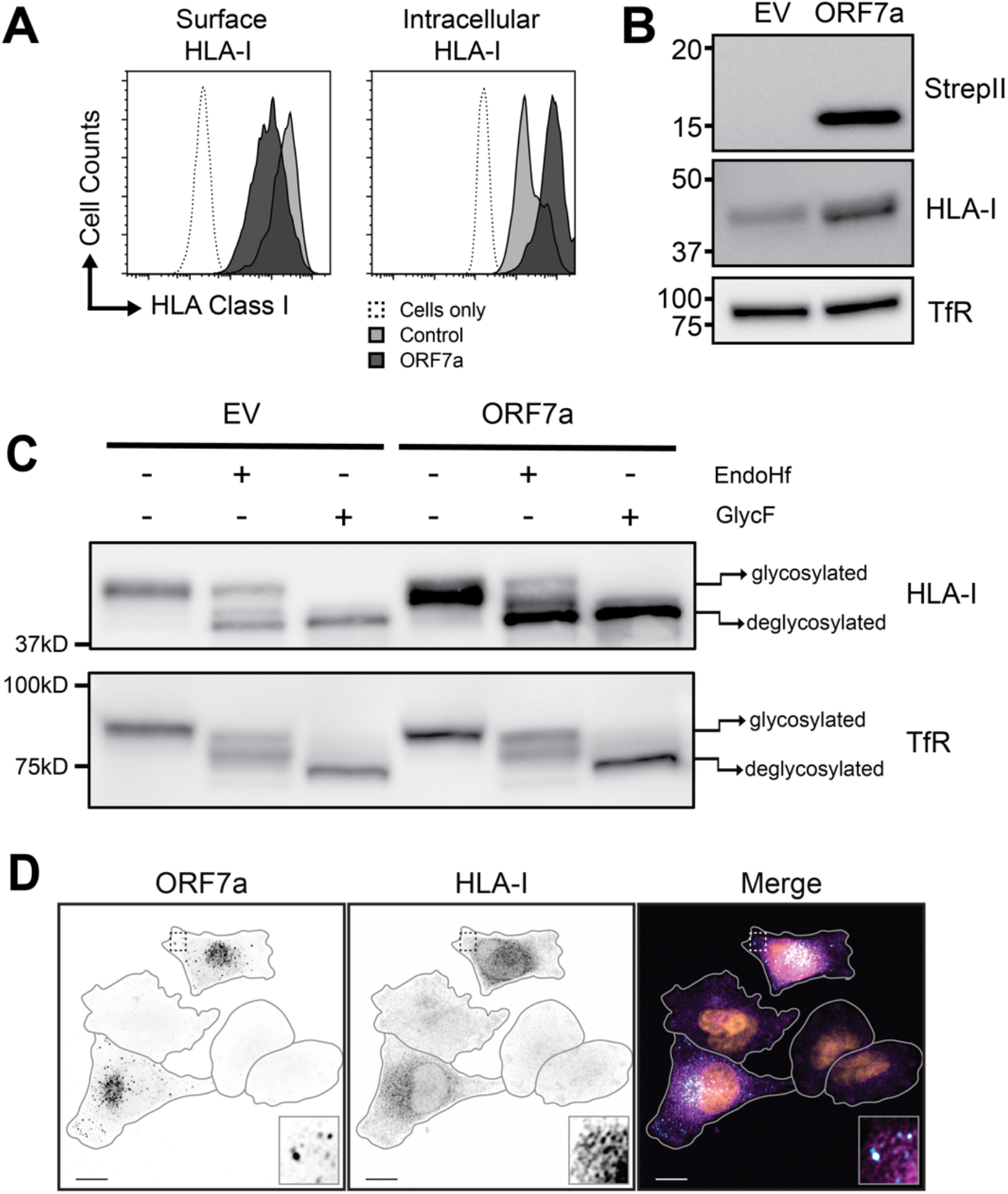
ORF7a induces intracellular HLA-I accumulation in the ER. (**A**) 293T cells transduced with ORF7a were subjected to surface and intracellular HLA-I stainings using the W6/32 antibody and assessed by flow cytometry at 7 dpi. The ORF7a vector co-expresses mAmetrine to allow discrimination between ORF7a-transduced (dark gray histograms, mAmetrine-positive) and non-transduced (light gray histograms, mAmetrine-negative) cells within the same culture. Unstained cells are indicated (open histograms). One representative experiment of at least 5 experiments is presented. (**B**) ORF7a, HLA-I and transferrin protein levels were assessed by immunoblotting of lysates from 293T cells transduced with an empty vector control (EV) or the SARS-CoV-2 ORF7a at 7 dpi; cells were selected to purity by puromycin prior to immunoblotting. (**C**) HLA-I and transferrin protein levels and glycosylation status were assessed by immunoblotting of 293T cells transduced with an empty control vector (EV) or ORF7a. Lysates were either left untreated or were incubated with Endo H_f_ or N-Glycosidase F (GlcF) prior to loading. (**D**) ORF7a and HLA-I localization were assessed by immunofluorescence in MelJuSo cells transduced with SARS-CoV-2 ORF7a. Nuclei were stained with DAPI. In the view, two ORF7a positive cells and three non-transduced cells are apparent. In the merged image, ORF7a signal is indicated in cyan, HLA-I signal in magenta, and DAPI signal in orange. The scale bar corresponds to 10 µm. Contours of cells are marked with a gray line and the insets show magnifications of the indicated region.

HLA class I HCs carry an N-glycan in their luminal/extracellular domain. Endoglycosidase H (EndoH)-treatment of lysates can provide information on the composition of the N-linked glycans of HLA-I HCs and, hence, their location in the cell. EndoH can only cleave immature N-linked glycans present on ER-resident glycoproteins, but not mature glycans present on proteins that have migrated to the Golgi and beyond. To assess where HLA-I molecules accumulate in ORF7a-transduced 293T cells, we incubated cell lysates with either EndoH or N-glycosidase. In control 293T cells, roughly half of the HLA-I molecules were ER-resident, as they were sensitive to EndoH-cleavage (Fig 6C, lane 2). ORF7a-expression induced upregulation of HLA class I (Fig 6C, lane 4), which was almost exclusively localized to the ER, as the products were sensitive to EndoH-activity resulting in deglycosylation of the majority of upregulated HLA-molecules (Fig 6C, compare lane 4 with lane 5). Treatment of samples with peptide N-glycanase-F (GlycF), which cleaves all glycans regardless of their maturation status, resulted in full cleavage of the N-glycan and provided a marker to visualize the mobility of deglycosylated HLA-I HCs (Figure 6C, Lane 6).

In agreement with the EndoH-treatment experiments, immunofluorescent imaging of MelJuSo cells transduced with ORF7a indicated that HLA-I accumulated in transduced cells in perinuclear compartments, including the ER and potentially the ER-Golgi intermediate compartment (ERGIC). Additionally, ORF7a-positive vesicles associated with HLA-I-positive structures. Our results suggest that ORF7a induces intracellular accumulation of HLA-I in the ER.

### ORF7a-mediated surface HLA-I downregulation is abrogated upon proteasome- and p97-inhibition, whereas intracellular HLA levels are unaffected

HLA class I downregulation by ER-localized viral inhibitors can be mediated by ER-associated protein degradation (ERAD). Dislocation of proteins from the ER back to the cytosol for proteasomal degradation is a hallmark in ERAD and p97, a cytosolic AAA-ATPase that is essential for ERAD, provides the driving force for this process. We next assessed whether the proteasome or p97 are involved in ORF7a-mediated HLA-I surface downregulation. For this, we treated control and ORF7a-transduced 293T cells with either the MG132 proteasome inhibitor or the CB-5083 p97 inhibitor (p97i) for four hours prior assessment of surface and intracellular levels of HLA class I (Fig. 7). As expected, in ORF7a-expressing cells and DMSO-treated ORF7a-expressing cells, we observed downregulation of surface HLA-I, accompanied by upregulation of intracellular HLA class I levels. Treatment with either MG132 or p97 inhibitor, however, abrogated ORF7a-mediated HLA class I surface downregulation without impacting intracellular HLA-I protein levels. Control cells transduced with the empty vector (EV) did not show regulation of intracellular nor surface HLA-I in MG132 and p97-inhibitor treated cells. Our results suggest that ORF7a-mediated HLA class I surface downregulation is dependent on p97 activity and the cellular proteasome system, possibly by engaging the cellular ERAD machinery.

**Figure 7.**
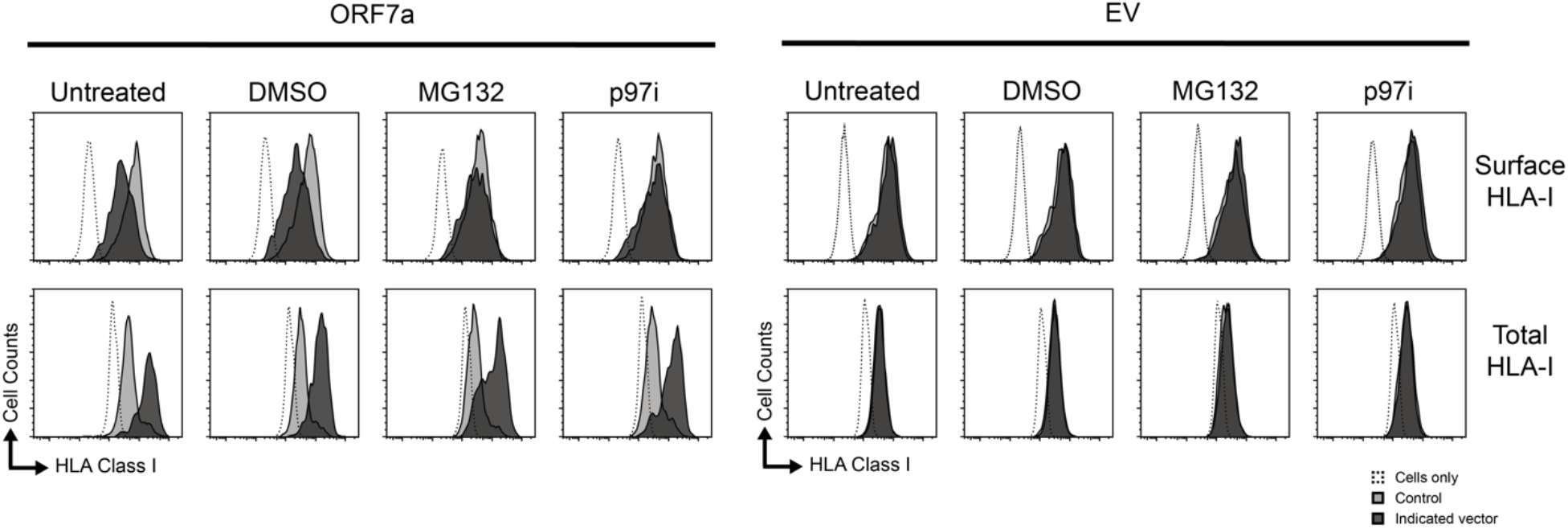
ORF7a-mediated surface HLA-I downregulation is abrogated upon proteasome- and p97-inhibition, whereas intracellular HLA-I levels are unaffected. 293T cells transduced with SARS-CoV-2 ORF7a or empty vector control (EV) were treated at 10 dpi with 20 µM MG132 or 4 µM CB-5083 p97-inhibitor for 4 hours. Subsequently, cells were harvested and subjected to surface and intracellular HLA-I stainings using the W6/32 antibody and analyzed by flow cytometry. The EV and ORF7a transduced cells co-express mAmetrine to allow discrimination between transduced (closed black histograms, mAmetrine-positive) and non-transduced (open histograms, mAmetrine-negative) cells within the same culture. One representative experiment of at least 5 experiments is presented.

### ORF7a associates with HLA class I molecules

ORF7a regulates HLA class I surface levels and co-localizes to the ER. We next asked whether ORF7a directly associates with HLA-I. We transduced 293T cells with StrepII-tagged ORF7a or empty vector and immunoprecipitated StrepII-tagged proteins from the lysate by means of Streptactin beads. Subsequent immunoblotting for ORF7a showed a strong enrichment for the protein (Fig 8, middle panel, lane 4), and co-precipitation of HLA class I was apparent (Fig 8, lower panel, lane 4). This co-precipitation was not observed in control cells transduced with the EV. Additionally, as control, the transferrin receptor (TfR) did not co-precipitate upon ORF7a pull down.

**Figure 8.**
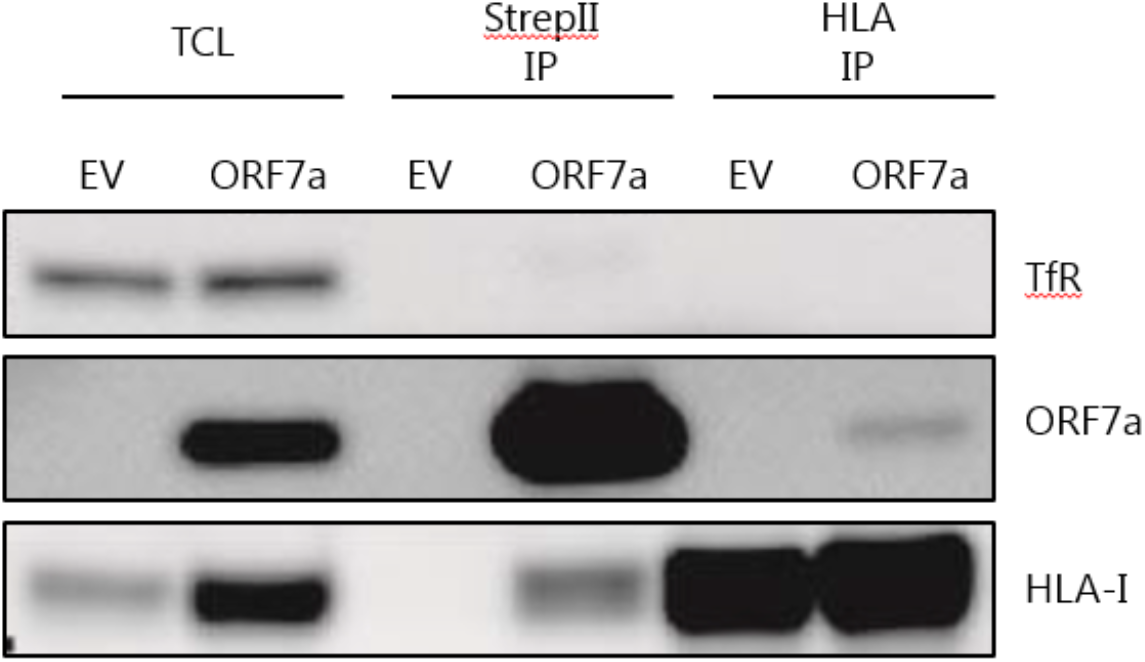
ORF7a interacts with HLA class I heavy chains. StrepII-tagged proteins were immunoprecipitated by using Streptactin beads from lysates of 293T cells transduced with EV or Strep-II-tagged ORF7a (StrepII IP: lanes 3 and 4). Furthermore, endogenous HLA class I molecules were immunoprecipitated from the same lysates by using Protein G Sepharose beads coupled to the HC10 anti-HLA class I antibody (HLA-I IP: lanes 5 and 6). Total cell lysates (TCL, lanes 1 and 2) and immunoprecipitation samples (lanes 3 through 6) were loaded on Western blot and stained for ORF7a, HLA class I, and TfR.

Also vice versa, ORF7a was specifically co-precipitated upon HLA class I pull down using the HC10 anti-HLA-I antibody. Thus, ORF7a is associated with HLA class I molecules in the cell, potentially via a direct interaction between the proteins. It is suggestive that the luminal domain of ORF7a interacts with the luminal domain of HLA-I HCs, as both are situated in the ER and mutations in the ORF7a luminal domain region (Fig 5) impact activity.

In summary, we show that ORF7a from SARS-CoV-2 downregulates HLA class I molecules from the surface of human cells. ORF7a interacts with HLA class I molecules, likely in the ER, resulting in increased intracellular HLA class I levels, yet abrogating HLA class I surface expression. ORF7a is dependent on its signal peptide sequence and transmembrane domain to mediate HLA class I surface downregulation, whereas the conserved ER-retrieval motif in its cytoplasmic tail is dispensable for this activity. Additionally, we show that ORF7a from SARS-CoV-1 does not direct surface HLA class I downregulation, but that a single amino acid substitution to the corresponding SARS-CoV-2 ORF7a sequence (T59F) yields a gain-of-function phenotype in which surface HLA class I molecules are downregulated. Importantly, HLA class I surface expression is impaired in SARS-CoV-2 infection models, which could partially be mediated by ORF7a activity. We hypothesize that SARS-CoV-2 can evade cytotoxic T lymphocytes immune responses towards infected cells via ORF7a-mediated interference with antigen presentation thereby evading immune surveillance. The ability of SARS-CoV-2 to interfere with its hosts adaptive immune system may contribute to the pathology of COVID-19.

## Discussion

Viral infections induce innate and adaptive immune responses in their host that inhibit virus replication and spread. As countermeasure, many viruses have evolved strategies to interfere with the antiviral immune response [19–21]. Here, we identified that ORF7a from SARS-CoV-2 can thwart the antigen presentation pathway by downregulating surface HLA class I surface expression, thereby potentially evading cytotoxic T cell responses targeted to virus-infected cells. HLA class I genes are highly polymorphic and may contribute to susceptibility and severity of COVID-19 [22–27]. Based on *in silico* analysis of HLA class I binding affinity for SARS-CoV-2 peptides, Nguyen *et al*. suggested that patients carrying the HLA-B*4601 genotype are more vulnerable to COVID-19, as this variant has a relatively low binding affinity for SARS-CoV-2 peptides [22]. This same genotype was associated with severe disease caused by SARS-CoV-1 in another study [28]. Furthermore, individuals with a HLA-A*01:01 and HLA-C*04:01 genotype have an increased risk to develop severe clinical course of COVID-19 [24, 25]. Kreutmair *et al*. employed NGS-based HLA class I typing in a patient cohort and integrated the data with single-cell immune profiling analysis to severe COVID-19-associated immune features [27]. They concluded that weak HLA class I binding, and thus impaired virus recognition, could partially drive the immunopathology of COVID-19. These results support earlier studies where patients with mild disease presented HLA class molecules with a higher theoretical capacity for binding SARS-CoV-2 peptides as compared to moderate and severe COVID-19 groups [29]. These studies illustrate that antigen presentation in general is important to limit SARS-CoV-2 infection. In line with this, SARS-CoV-2 can subvert CD8^+^ T cell surveillance through point mutations in HLA-I-restricted viral epitopes [30]. Hence, interference in antigen presentation by SARS-CoV-2 could be advantageous for the virus to escape from its host’s immune system. Besides by selecting for mutations in HLA-I viral epitopes, evasion can be mediated by viral gene-products that directly target antigen presentation, such as ORF7a.

SARS-CoV-2 ORF7a is a transmembrane protein which localizes to the Golgi/ERGIC [31–34]. ORF7a has a five amino acid cytosolic tail containing a dilysine (KRKTE) endoplasmic reticulum (ER) retrieval signal (ERRS). This motif is common for Golgi residing proteins and mediates protein trafficking to the ER-Golgi intermediate compartment (ERGIC) [14,33,35,36]. Mutation of the cytosolic tail of ORF7a (from KRKTE to KAAAA) did not abrogate HLA class I downregulation from the cell surface, suggesting that the motif is not required for this activity. However, studies on SARS-CoV-1 ORF7a suggest that additional residues in the transmembrane domain impact protein trafficking [33]. Indeed, by deleting the entire transmembrane domain and the cytosolic tail, ORF7a-mediated HLA class I surface downregulation was fully impaired, likely caused by disturbed localization of the protein and loss of its ability to reside at the ER/Golgi membrane. In line with this, a SARS-CoV-2 isolate with an 115 nt deletion in the ORF7a gene encompassing the transmembrane region and ERRS motif, displayed ORF7a expression in the cytosol instead of the perinuclear region [32]. Both SARS-CoV-1 ORF7a and SARS-CoV-2 ORF7a carry a 15 amino acid signal peptide at their N-terminus, which is cleaved from the native ORF7a upon entry into the ER [33]. Unexpectedly, however, replacement of the SARS-CoV-2 ORF7a signal peptide sequence with that of the CD8A protein abrogated the ability to downregulate HLA class I from the cell surface. This suggests that the ORF7a signal peptide, besides guiding the protein into the secretory pathway during translation, is involved in the ORF7a-mediated downregulation of surface HLA-I.

BST-2 (tetherin) restricts the release of viruses from cells and SARS-CoV-1 ORF7a has been shown to inhibit BST-2 by blocking its glycosylation [37]. The same has been shown for SARS-CoV-2 ORF7a [31]. A preprint showed that the loss of tetherin increased virus titers and spread of SARS-CoV-2, but that this effect was mediated by the viral spike protein, whereas SARS-CoV-2 ORF7a induced Golgi fragmentation [38]. The Golgi apparatus is crucial for HLA class I transport to the cell surface and it has been shown for various viruses, including SARS-CoV-1 and SARS-CoV-2, that infection can result in Golgi fragmentation [38–42]. Infection with Orf virus for example results in Golgi fragmentation and subsequent impairment of intracellular MHC class I transport [39]. ORF7a-mediated Golgi fragmentation may contribute to decreased HLA-I surface levels through impaired transport of heavy chains through the secretory pathway.

We show that ORF7a specifically downregulates HLA-I from the cell surface, as other cell-surface receptors that migrate through the secretory pathway are unaffected by the viral gene product. Also, our data suggests that ORF7a is localized to perinuclear vesicles, potentially associated with the ER; that the protein associates with HLA class I heavy chains; and that ORF7a expression results in accumulation of HLA-I heavy chains in the ER. These data suggest that ORF7a specifically binds to HLA-I HCs in the ER, either retaining HLA class I HC molecules, or interfering with their migration to the Golgi network for subsequent presentation to the cell surface. It is probable that a region in the ER luminal domain of ORF7a mediates the HLA-I HC binding, as we showed that SARS-CoV-1 ORF7a mutations in the luminal domain resulted in a gain-of-function of HLA-I downregulation. Upon inhibition of the proteasome or p97, ORF7a-mediated surface HLA-I downregulation was impaired. It is conceivable that blockage of HLA class I degradation via the proteasome in the face of ORF7a expression, results in enhanced HC accumulation in the ER thereby overloading ORF7a’s ability to engage HLA-I molecules, ultimately resulting in re-appearance of HLA-I expression on the cell surface. Alternatively, ORF7a may also engage the ERAD machinery directly to induce surface HLA-I downregulation, as has been observed for viral gene products of other viruses [43]. For example, HCMV US2 and US11 encode for ER transmembrane proteins that mediate HLA-I degradation by engaging the ERAD machinery [44, 45]. US2 and US11 bind to HLA-I HCs in the ER to promote their downregulation in concert with multiple host factors such as the E3 ubiquitin ligase TRC8 [46], the AAA-ATPase p97 [47] for US2 and the E3 ubiquitin ligase TMEM129 [48, 49], p97, Ufd1 and Npl4 [50] for US11. ERAD and p97 are responsible for the dislocation of proteins from the ER back to the cytosol for subsequent proteasomal degradation and could be involved in the ORF7a-driven surface downregulation of HLA-I as well. However, whereas US2 and US11 induce degradation of HLA-I HCs from the cell, ORF7a-expressing 293T and MelJuSo cells show increased intracellular HLA-I levels in conjunction with reduced cell-surface levels. Hence, ERAD-mediated class I degradation alone does not explain the observed phenotype. It is possible that ORF7a-mediated HLA-I surface downregulation and ORF7a-mediated intracellular accumulation are the result of two separate mechanisms. Indeed, our preliminary observations showed that, in Huh7 cells, ORF7a-mediated surface HLA-I downregulation occurs in the absence of intracellular HLA-I accumulation (data not shown), suggesting that the processes may be decoupled and are cell-type specific. The molecular mechanism of HLA-I surface downregulation by SARS-CoV-2 ORF7a remains unknown and awaits further clarification.

Besides ORF7a, multiple studies have identified SARS-CoV-2 gene products to interfere with MHC-I expression. A recent preprint showed that several SARS-CoV-2 gene products have the capacity to downregulate MHC-I molecules, including the viral E, M, ORF7a, ORF7b, and ORF8 proteins [51]. In this study, no additional experiments were performed to address the nature of MHC-I regulation. Zhang *et al.* reported that SARS-CoV-2 ORF8 selectively targets HLA class I molecules for lysosomal degradation in an autophagy-dependent manner [13]. Contrary to our findings, the authors reported that ORF7a did not result in surface HLA class I downregulation in HEK-293T cells. Another recent study identified SARS-CoV-2 ORF6 to suppress the HLA-I transactivator NLRC5, thereby inhibiting the induction of HLA-I gene expression, without impacting steady-state HLA-I expression levels [52]. Also here, the authors did not observe an effect of ORF8 on HLA-I expression, whereas ORF7a was not included in their experiments. In our study, we do not find experimental evidence for a role of either ORF6 nor ORF8 on steady-state HLA class I surface expression levels, which agrees with Yoo *et al.* [52], but not with Zhang *et al*. [13]. It is not clear why the inhibitory effect of ORF8 on HLA class I expression was not observed in our study; potentially, ORF8 activity can only be observed under specific experimental conditions. In line with this reasoning, we observed a transient effect of SARS-CoV-2 ORF7a mediated HLA-I downregulation (data not shown), and the potency of downregulation varied between experiments. Therefore, ORF7a-mediated HLA-I regulation may be overlooked by others. Despite these different findings, it is remarkable that SARS-CoV-2 has evolved multiple strategies to thwart with HLA class I antigen presentation. To assess the specific contribution of ORF7a on HLA-I downregulation during infection, we used a recombinant SARS-CoV-2 strain in which the ORF7a gene was substituted with the GFP gene [9]. This virus however, induced equivalent HLA-I surface downregulation as compared to the parental ORF7a-encoding SARS-CoV-2 strain in both Vero E6 and A549-ACE2-TMPRSS2 cells (Supplemental Fig.1). It is likely that abrogation of the ORF7a-mediated HLA-I downregulation alone is not sufficient to contribute to a detectable increase of surface HLA-I levels, as multiple viral gene products that target HLA-I are still functionally expressed (ORF6, ORF8, E, M). If, and to what extent, each of these genes contribute to the inhibition of cytotoxic T cell responses *in vivo* remains to be determined.

SARS-CoV-1 and SARS-CoV-2 ORF7a have a conserved immunoglobulin-fold carrying an integrin binding site and the protein displays structural homology to ICAM-1 [53]. ICAM-1 is a ligand for the T cell integrin receptor LFA-1 [54] and their engagement is important for T cell activation and motility [55, 56]. Various deletions have been reported in the ORF7a gene, some resulting in ORF7a/7b or ORF7a/8 fusion products [32,57–59]. In these strains, amino acid E26 is maintained, which is a position in the ORF7a Ig fold that harbours the integrin binding region for LFA-1 (and Mac-1) [53]. Interestingly, both ORF7a and ORF8 carry an Ig-fold and share distant nucleotide sequence similarity [60–64], suggesting that they are paralogs that could share similar immune antagonistic functions. Neither ORF7a nor ORF8 are present in members of the gamma- or deltacoronaviruses and, most surprisingly, not in MERS [60].

By replacing several amino acids of the luminal domain of SARS-CoV-1 ORF7a with that of the corresponding SARS-CoV-2 ORF7a sequence, SARS-CoV-1 ORF7a gained the ability to downregulate surface HLA-I expression levels. Intriguingly, only a single amino acid substitution (T59F) was responsible for the gain-of-function phenotype. This suggests that the F59 position of SARS-CoV-2 ORF7a is of key importance for ORF7a activity. Indeed, protein sequence alignment of multiple sarbecovirus ORF7a sequences indicated that the 59 position has significant variation. Where multiple sarbecoviruses ORF7a proteins carry a bulky hydrophobic phenylalanine (F) or isoleucine (I) at this position, other sarbecoviruses have a small polar amino acid at this site (T: Threonine, or S: Serine). Although we have not assessed whether ORF7a from additional sarbecoviruses can also abrogate surface MHC class I expression, it is likely that this ability is shared with other members of the family. It is striking that a single amino acid chance in ORF7a can direct such a fundamental change in phenotype.

In conclusion, our data shows that the SARS-CoV-2 accessory protein ORF7a directly associates with HLA-I and inhibits HLA-I surface expression levels in human cells. By abrogating HLA class I antigen presentation, ORF7a may facilitate the evasion of cytotoxic T cell responses thereby acting pro-viral. Future therapeutical interventions by targeting ORF7a may lead to improved clearance of SARS-CoV-2 infected cells thereby limiting infection and spread.

### Note added prior submission

In the process of submitting this paper, two preprint were published in which SARS-CoV-2 ORF7a was also identified to regulate HLA-I expression [65, 66]. The data presented in these preprints are largely in agreement with our findings.

## Acknowledgements

We would like to thank the members of the Wiertz and Lebbink lab for conducive discussions. This work was supported by grants from the European Commission under the Horizon2020 program H2020 MSCA-ITN GA 675278 EDGE to EJHJW and RJL Health∼Holland (grant number LSHM20058) to JMB, MN, and RJL; and Deutsche Forschungsgemeinschaft (DFG) grant GK1949/2 to ID. Further support came from the European Research Council (ERC Consolidator Grant 819219 to L.C.K.) and the Centre for Living Technologies, a part of the Alliance TU/e, WUR, UU, UMC Utrecht (www.ewuu.nl).

## Author contributions

SZ, HdB, PP, and RJL conceived and designed the experiments. HdB and PP performed initial experiments to identify ORF7a as inhibitor of HLA-I expression. MvG and WN performed the immunofluorescence experiments. SZ, HdB, and AE generated plasmids and reagents. YL and ID contributed reagents. PP and MvG performed live SARS-CoV-2 experiments. AE performed the proteasome/p97 inhibitor experiments, IBB performed all biochemical experiments. LdV assisted with flow cytometry stainings. SZ performed all other experiments. LCK, JMB, MN, EJHJW, and RJL provided support, reagents, and funding. SZ, HdB, and RJL wrote the paper with input from all the other authors.

## Notes

### Competing Interest Statement

The authors have declared no competing interest.

## References

1. Cui J, Li F, Shi ZL: Origin and evolution of pathogenic coronaviruses. Nat Rev Microbiol 2019, 17:181–192.

2. Fung TS, Liu DX: Human Coronavirus: Host-Pathogen Interaction. Annu Rev Microbiol 2019, 73:529–557.

3. Hartenian E, Nandakumar D, Lari A, Ly M, Tucker JM, Glaunsinger BA: The molecular virology of coronaviruses. J Biol Chem 2020, 295:12910–12934.

4. Robson F, Khan KS, Le TK, Paris C, Demirbag S, Barfuss P, Rocchi P, Ng WL: Coronavirus RNA Proofreading: Molecular Basis and Therapeutic Targeting. Mol Cell 2020, 79:710–727.

5. Harrison AG, Lin T, Wang P: Mechanisms of SARS-CoV-2 Transmission and Pathogenesis. Trends Immunol 2020, 41:1100–1115.

6. V’Kovski P, Kratzel A, Steiner S, Stalder H, Thiel V: Coronavirus biology and replication: implications for SARS-CoV-2. Nat Rev Microbiol 2020.

7. Golstein P, Griffiths GM: An early history of T cell-mediated cytotoxicity. Nat Rev Immunol 2018, 18:527–535.

8. Praest P, de Buhr H, Wiertz E: A Flow Cytometry-Based Approach to Unravel Viral Interference with the MHC Class I Antigen Processing and Presentation Pathway. Methods Mol Biol 2019, 1988:187–198.

9. Thi Nhu Thao T, Labroussaa F, Ebert N, V’Kovski P, Stalder H, Portmann J, Kelly J, Steiner S, Holwerda M, Kratzel A, et al.: Rapid reconstruction of SARS-CoV-2 using a synthetic genomics platform. Nature 2020, 582:561–565.

10. Gordon DE, Hiatt J, Bouhaddou M, Rezelj VV, Ulferts S, Braberg H, Jureka AS, Obernier K, Guo JZ, Batra J, et al.: Comparative host-coronavirus protein interaction networks reveal pan-viral disease mechanisms. Science 2020, 370.

11. van de Weijer ML, van Muijlwijk GH, Visser LJ, Costa AI, Wiertz EJ, Lebbink RJ: The E3 Ubiquitin Ligase TMEM129 Is a Tri-Spanning Transmembrane Protein. Viruses 2016, 8.

12. Redondo N, Zaldivar-Lopez S, Garrido JJ, Montoya M: SARS-CoV-2 Accessory Proteins in Viral Pathogenesis: Knowns and Unknowns. Front Immunol 2021, 12:708264.

13. Zhang Y, Chen Y, Li Y, Huang F, Luo B, Yuan Y, Xia B, Ma X, Yang T, Yu F, et al.: The ORF8 protein of SARS-CoV-2 mediates immune evasion through down-regulating MHC-I. Proc Natl Acad Sci U S A 2021, 118.

14. Fielding BC, Tan YJ, Shuo S, Tan TH, Ooi EE, Lim SG, Hong W, Goh PY: Characterization of a unique group-specific protein (U122) of the severe acute respiratory syndrome coronavirus. J Virol 2004, 78:7311–7318.

15. Holcomb D, Alexaki A, Hernandez N, Hunt R, Laurie K, Kames J, Hamasaki-Katagiri N, Komar AA, DiCuccio M, Kimchi-Sarfaty C: Gene variants of coagulation related proteins that interact with SARS-CoV-2. PLoS Comput Biol 2021, 17:e1008805.

16. Tan YJ, Fielding BC, Goh PY, Shen S, Tan TH, Lim SG, Hong W: Overexpression of 7a, a protein specifically encoded by the severe acute respiratory syndrome coronavirus, induces apoptosis via a caspase-dependent pathway. J Virol 2004, 78:14043–14047.

17. Schaecher SR, Touchette E, Schriewer J, Buller RM, Pekosz A: Severe acute respiratory syndrome coronavirus gene 7 products contribute to virus-induced apoptosis. J Virol 2007, 81:11054–11068.

18. Tan YX, Tan TH, Lee MJ, Tham PY, Gunalan V, Druce J, Birch C, Catton M, Fu NY, Yu VC, et al.: Induction of apoptosis by the severe acute respiratory syndrome coronavirus 7a protein is dependent on its interaction with the Bcl-XL protein. J Virol 2007, 81:6346–6355.

19. Schuren AB, Costa AI, Wiertz EJ: Recent advances in viral evasion of the MHC Class I processing pathway. Curr Opin Immunol 2016, 40:43–50.

20. van de Weijer ML, Luteijn RD, Wiertz EJ: Viral immune evasion: Lessons in MHC class I antigen presentation. Semin Immunol 2015, 27:125–137.

21. Kasuga Y, Zhu B, Jang KJ, Yoo JS: Innate immune sensing of coronavirus and viral evasion strategies. Exp Mol Med 2021, 53:723–736.

22. Nguyen A, David JK, Maden SK, Wood MA, Weeder BR, Nellore A, Thompson RF: Human Leukocyte Antigen Susceptibility Map for Severe Acute Respiratory Syndrome Coronavirus 2. J Virol 2020, 94.

23. Migliorini F, Torsiello E, Spiezia F, Oliva F, Tingart M, Maffulli N: Association between HLA genotypes and COVID-19 susceptibility, severity and progression: a comprehensive review of the literature. Eur J Med Res 2021, 26:84.

24. Shkurnikov M, Nersisyan S, Jankevic T, Galatenko A, Gordeev I, Vechorko V, Tonevitsky A: Association of HLA Class I Genotypes With Severity of Coronavirus Disease-19. Front Immunol 2021, 12:641900.

25. Weiner J, Suwalski P, Holtgrewe M, Rakitko A, Thibeault C, Muller M, Patriki D, Quedenau C, Kruger U, Ilinsky V, et al.: Increased risk of severe clinical course of COVID-19 in carriers of HLA-C*04:01. EClinicalMedicine 2021, 40:101099.

26. Douillard V, Castelli EC, Mack SJ, Hollenbach JA, Gourraud PA, Vince N, Limou S, Covid, Hla, Immunogenetics C, et al.: Current HLA Investigations on SARS-CoV-2 and Perspectives. Front Genet 2021, 12:774922.

27. Kreutmair S, Unger S, Nunez NG, Ingelfinger F, Alberti C, De Feo D, Krishnarajah S, Kauffmann M, Friebel E, Babaei S, et al.: Distinct immunological signatures discriminate severe COVID-19 from non-SARS-CoV-2-driven critical pneumonia. Immunity 2021, 54:1578–1593 e1575.

28. Lin M, Tseng HK, Trejaut JA, Lee HL, Loo JH, Chu CC, Chen PJ, Su YW, Lim KH, Tsai ZU, et al.: Association of HLA class I with severe acute respiratory syndrome coronavirus infection. BMC Med Genet 2003, 4:9.

29. Iturrieta-Zuazo I, Rita CG, Garcia-Soidan A, de Malet Pintos-Fonseca A, Alonso-Alarcon N, Pariente-Rodriguez R, Tejeda-Velarde A, Serrano-Villar S, Castaner-Alabau JL, Nieto-Ganan I: Possible role of HLA class-I genotype in SARS-CoV-2 infection and progression: A pilot study in a cohort of Covid-19 Spanish patients. Clin Immunol 2020, 219:108572.

30. Agerer B, Koblischke M, Gudipati V, Montano-Gutierrez LF, Smyth M, Popa A, Genger JW, Endler L, Florian DM, Muhlgrabner V, et al.: SARS-CoV-2 mutations in MHC-I-restricted epitopes evade CD8(+) T cell responses. Sci Immunol 2021, 6.

31. Martin-Sancho L, Lewinski MK, Pache L, Stoneham CA, Yin X, Becker ME, Pratt D, Churas C, Rosenthal SB, Liu S, et al.: Functional landscape of SARS-CoV-2 cellular restriction. Mol Cell 2021.

32. Nemudryi A, Nemudraia A, Wiegand T, Nichols J, Snyder DT, Hedges JF, Cicha C, Lee H, Vanderwood KK, Bimczok D, et al.: SARS-CoV-2 genomic surveillance identifies naturally occurring truncation of ORF7a that limits immune suppression. Cell Rep 2021:109197.

33. Nelson CA, Pekosz A, Lee CA, Diamond MS, Fremont DH: Structure and intracellular targeting of the SARS-coronavirus Orf7a accessory protein. Structure 2005, 13:75–85.

34. Zhang J, Cruz-Cosme R, Zhuang MW, Liu D, Liu Y, Teng S, Wang PH, Tang Q: A systemic and molecular study of subcellular localization of SARS-CoV-2 proteins. Signal Transduct Target Ther 2020, 5:269.

35. Giraudo CG, Maccioni HJ: Endoplasmic reticulum export of glycosyltransferases depends on interaction of a cytoplasmic dibasic motif with Sar1. Mol Biol Cell 2003, 14:3753–3766.

36. Ujike M, Huang C, Shirato K, Makino S, Taguchi F: The contribution of the cytoplasmic retrieval signal of severe acute respiratory syndrome coronavirus to intracellular accumulation of S proteins and incorporation of S protein into virus-like particles. J Gen Virol 2016, 97:1853–1864.

37. Taylor JK, Coleman CM, Postel S, Sisk JM, Bernbaum JG, Venkataraman T, Sundberg EJ, Frieman MB: Severe Acute Respiratory Syndrome Coronavirus ORF7a Inhibits Bone Marrow Stromal Antigen 2 Virion Tethering through a Novel Mechanism of Glycosylation Interference. J Virol 2015, 89:11820–11833.

38. Stewart H, Johansen K, McGovern N, Palmulli R, Carnell G, Heeney J, Okkenhaug K, Firth A, Peden A, Edgar J: SARS-CoV-2 spike downregulates tetherin to enhance viral spread. bioRxiv 2021:2021.2001.2006.425396.

39. Rohde J, Emschermann F, Knittler MR, Rziha HJ: Orf virus interferes with MHC class I surface expression by targeting vesicular transport and Golgi. BMC Vet Res 2012, 8:114.

40. Hansen MD, Johnsen IB, Stiberg KA, Sherstova T, Wakita T, Richard GM, Kandasamy RK, Meurs EF, Anthonsen MW: Hepatitis C virus triggers Golgi fragmentation and autophagy through the immunity-related GTPase M. Proc Natl Acad Sci U S A 2017, 114:E3462–E3471.

41. Mousnier A, Swieboda D, Pinto A, Guedan A, Rogers AV, Walton R, Johnston SL, Solari R: Human rhinovirus 16 causes Golgi apparatus fragmentation without blocking protein secretion. J Virol 2014, 88:11671–11685.

42. Freundt EC, Yu L, Goldsmith CS, Welsh S, Cheng A, Yount B, Liu W, Frieman MB, Buchholz UJ, Screaton GR, et al.: The open reading frame 3a protein of severe acute respiratory syndrome-associated coronavirus promotes membrane rearrangement and cell death. J Virol 2010, 84:1097–1109.

43. Morito D, Nagata K: Pathogenic Hijacking of ER-Associated Degradation: Is ERAD Flexible? Mol Cell 2015, 59:335–344.

44. Wiertz EJ, Tortorella D, Bogyo M, Yu J, Mothes W, Jones TR, Rapoport TA, Ploegh HL: Sec61-mediated transfer of a membrane protein from the endoplasmic reticulum to the proteasome for destruction. Nature 1996, 384:432–438.

45. Wiertz EJ, Jones TR, Sun L, Bogyo M, Geuze HJ, Ploegh HL: The human cytomegalovirus US11 gene product dislocates MHC class I heavy chains from the endoplasmic reticulum to the cytosol. Cell 1996, 84:769–779.

46. Stagg HR, Thomas M, van den Boomen D, Wiertz EJ, Drabkin HA, Gemmill RM, Lehner PJ: The TRC8 E3 ligase ubiquitinates MHC class I molecules before dislocation from the ER. J Cell Biol 2009, 186:685–692.

47. Soetandyo N, Ye Y: The p97 ATPase dislocates MHC class I heavy chain in US2-expressing cells via a Ufd1-Npl4-independent mechanism. J Biol Chem 2010, 285:32352–32359.

48. van de Weijer ML, Bassik MC, Luteijn RD, Voorburg CM, Lohuis MA, Kremmer E, Hoeben RC, LeProust EM, Chen S, Hoelen H, et al.: A high-coverage shRNA screen identifies TMEM129 as an E3 ligase involved in ER-associated protein degradation. Nat Commun 2014, 5:3832.

49. van den Boomen DJ, Timms RT, Grice GL, Stagg HR, Skodt K, Dougan G, Nathan JA, Lehner PJ: TMEM129 is a Derlin-1 associated ERAD E3 ligase essential for virus-induced degradation of MHC-I. Proc Natl Acad Sci U S A 2014, 111:11425–11430.

50. Ye Y, Meyer HH, Rapoport TA: The AAA ATPase Cdc48/p97 and its partners transport proteins from the ER into the cytosol. Nature 2001, 414:652–656.

51. Moriyama M, Lucas C, Monteiro VS, Yale S-C-GSI, Iwasaki A: SARS-CoV-2 variants do not evolve to promote further escape from MHC-I recognition. bioRxiv 2022.

52. Yoo JS, Sasaki M, Cho SX, Kasuga Y, Zhu B, Ouda R, Orba Y, de Figueiredo P, Sawa H, Kobayashi KS: SARS-CoV-2 inhibits induction of the MHC class I pathway by targeting the STAT1-IRF1-NLRC5 axis. Nat Commun 2021, 12:6602.

53. Nizamudeen ZA, Xu ER, Karthik V, Halawa M, Arkill KP, Jackson AM, Bates DO, Emsley J: Structural assessment of SARS-CoV2 accessory protein ORF7a predicts LFA-1 and Mac-1 binding potential. Biosci Rep 2021, 41.

54. Hanel K, Willbold D: SARS-CoV accessory protein 7a directly interacts with human LFA-1. Biol Chem 2007, 388:1325–1332.

55. Walling BL, Kim M: LFA-1 in T Cell Migration and Differentiation. Front Immunol 2018, 9:952.

56. Bui TM, Wiesolek HL, Sumagin R: ICAM-1: A master regulator of cellular responses in inflammation, injury resolution, and tumorigenesis. J Leukoc Biol 2020, 108:787–799.

57. Holland LA, Kaelin EA, Maqsood R, Estifanos B, Wu LI, Varsani A, Halden RU, Hogue BG, Scotch M, Lim ES: An 81-Nucleotide Deletion in SARS-CoV-2 ORF7a Identified from Sentinel Surveillance in Arizona (January to March 2020). J Virol 2020, 94.

58. Addetia A, Xie H, Roychoudhury P, Shrestha L, Loprieno M, Huang ML, Jerome KR, Greninger AL: Identification of multiple large deletions in ORF7a resulting in in-frame gene fusions in clinical SARS-CoV-2 isolates. J Clin Virol 2020, 129:104523.

59. Rosenthal SH, Kagan RM, Gerasimova A, Anderson B, Grover D, Hua M, Liu Y, Owen R, Lacbawan F: Identification of eight SARS-CoV-2 ORF7a deletion variants in 2,726 clinical specimens. bioRxiv 2020:2020.2012.2010.418855.

60. Tan Y, Schneider T, Leong M, Aravind L, Zhang D: Novel Immunoglobulin Domain Proteins Provide Insights into Evolution and Pathogenesis of SARS-CoV-2-Related Viruses. mBio 2020, 11.

61. Neches RY, Kyrpides NC, Ouzounis CA: Atypical Divergence of SARS-CoV-2 Orf8 from Orf7a within the Coronavirus Lineage Suggests Potential Stealthy Viral Strategies in Immune Evasion. mBio 2021, 12.

62. Young BE, Fong SW, Chan YH, Mak TM, Ang LW, Anderson DE, Lee CY, Amrun SN, Lee B, Goh YS, et al.: Effects of a major deletion in the SARS-CoV-2 genome on the severity of infection and the inflammatory response: an observational cohort study. Lancet 2020, 396:603–611.

63. Zinzula L: Lost in deletion: The enigmatic ORF8 protein of SARS-CoV-2. Biochem Biophys Res Commun 2021, 538:116–124.

64. Flower TG, Buffalo CZ, Hooy RM, Allaire M, Ren X, Hurley JH: Structure of SARS-CoV-2 ORF8, a rapidly evolving coronavirus protein implicated in immune evasion. bioRxiv 2020.

65. Arshad N, Laurent-Rolle M, Ahmed WS, Hsu JC, Mitchell SM, Pawlak J, Sengupta D, Biswas KH, Cresswell P: SARS-CoV-2 accessory proteins ORF7a and ORF3a use distinct mechanisms to downregulate MHC-I surface expression. bioRxiv 2022.

66. Zhang F, Zang T, Stevenson EM, Lei X, Copertino DC, Mota TM, Boucau J, Garcia-Beltran WF, Jones B, Bieniasz PD: Inhibition of major histocompatibility complex-I antigen presentation by sarbecovirus ORF7a proteins. bioRxiv 2022.

