## Supplemental Figure 1 for "The SARS-CoV-2 accessory factor ORF7a downregulates MHC class I surface expression"

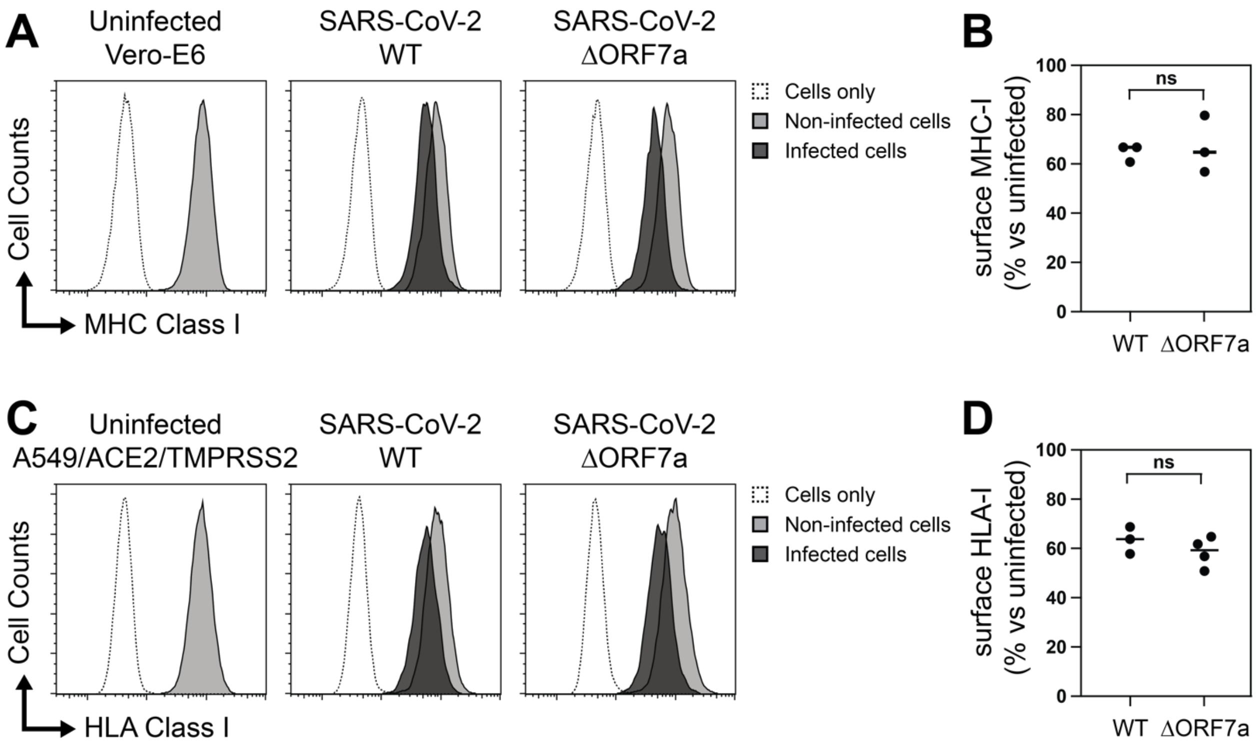


**Supplemental Figure 1. Deletion of ORF7a from SARS-CoV-2 does not abrogate surface MHC class I downregulation in Vero E6 and A549 cells.**

(**A**) Vero E6 cells were infected with a recombinant SARS-CoV-2 virus [1] encoding the GFP gene at the ORF7a locus (ΔORF7a) and the parental SARS-CoV-2 isolate harbouring an intact ORF7a gene at an MOI of 0.05. Cells were subsequently analyzed for surface MHC class I levels by flow cytometry at 2dpi. Spike antibody stains were included to allow discrimination between SARS-CoV-2 infected (infected, black filled histograms) and non-infected cells present in the same culture (gray histograms). Unstained cells are indicated (open histograms). One representative experiment of three is shown. (**B**) Quantification of experiments as described in A). Surface MHC-I levels of cells infected with the indicated viruses are normalized to uninfected Spike-negative cells in the same culture. (**C)** same as in A), yet A549-ACE2-TMPRSS2 cells were infected with the indicated viruses at an MOI of 0.5. (**D**) Quantification of experiments as described in C). Surface HLA-I levels of cells infected with the indicated viruses are normalized to uninfected Spike-negative cells in the same culture.

1. Thi Nhu Thao T, Labroussaa F, Ebert N, V'Kovski P, Stalder H, Portmann J, Kelly J, Steiner S, Holwerda M, Kratzel A, et al.: **Rapid reconstruction of SARS-CoV-2 using a synthetic genomics platform**. *Nature* 2020, **582**:561-565.
